# Effect of channel density, inverse solutions and connectivity measures on EEG resting-state networks: a simulation study

**DOI:** 10.1101/2022.06.01.494301

**Authors:** Sahar Allouch, Aya Kabbara, Joan Duprez, Mohamad Khalil, Julien Modolo, Mahmoud Hassan

## Abstract

Along with the study of brain activity evoked by external stimuli, the past two decades witnessed an increased interest in characterizing the spontaneous brain activity occurring during resting conditions. The identification of the connectivity patterns in this so-called “resting-state” has been the subject of a great number of electrophysiology-based studies, using the Electro/Magneto-Encephalography (EEG/MEG) source connectivity method. However, no consensus has been reached yet regarding a unified (if possible) analysis pipeline, and several involved parameters and methods require cautious tuning. This is particularly challenging when different choices induce significant discrepancy in results and drawn conclusions, thereby hindering reproducibility of neuroimaging research. Hence, our objective in this study was to evaluate some of the parameters related to the EEG source connectivity analysis and shed light on their implications on the accuracy of the resulting networks. We simulated, using neural mass models, EEG data corresponding to two of the resting-state networks (RSNs), namely the default mode network (DMN) and the dorsal attentional network (DAN). We investigated the impact of five channel densities (19, 32, 64, 128, 256), three inverse solutions (weighted minimum norm estimate (wMNE), exact low resolution brain electromagnetic tomography (eLORETA), and linearly constrained minimum variance (LCMV) beamforming) and four functional connectivity measures (phase-locking value (PLV), phase-lag index (PLI), and amplitude envelope correlation (AEC) with and without source leakage correction), on the correspondence between reconstructed and reference networks. We showed that, with different analytical choices, a high variability is present in the results. More specifically, our results show that a higher number of EEG channels significantly increased the accuracy of the reconstructed networks. Additionally, our results showed a significant variability in the performance of the tested inverse solutions and connectivity measures. In our specific simulation context, eLORETA and wMNE combined with AEC computed between orthogonalized time series exhibited the highest performance in terms of similarity between reconstructed and reference connectivity matrices. Results were similar for both DMN and DAN. We believe that this work could be useful for the field of electrophysiology connectomics, by shedding light on the challenge of analytical variability and its consequences on the reproducibility of neuroimaging studies.

## 1. Introduction

Over the past two decades, there has been a considerable growth in the number of studies investigating human brain activity at rest (Raichle et al. 2001; Raichle and Snyder 2007; van den Heuvel, Mandl, and Pol 2008; Damoiseaux et al. 2006). Characterizing synchronous activity across spatially distributed regions has revealed consistent patterns of brain connectivity in the absence of a goal-directed task, with a preserved consistency across subjects as well as across different neuroimaging modalities (Damoiseaux et al. 2006; Brookes et al. 2011; Francesco de Pasquale et al. 2012; Kabbara, Paban, and Hassan 2021). Interestingly, an accurate identification of the so-called resting-state networks (RSNs) has been found to be primordial for both cognitive (Keller et al. 2015; Jockwitz et al. 2017; Shen et al. 2018; Alavash et al. 2015) and clinical studies (Sheline and Raichle 2013; Kabbara et al. 2018; Hassan, Chaton, et al. 2017; Gratton et al. 2018; Jimenez et al. 2019).

Functional RSNs can be mapped, non-invasively, using either functional magnetic resonance imaging (fMRI) or magneto/electro-encephalography (M/EEG). The main advantage of EEG, as compared to fMRI, which is more established in resting-state studies, is its ability to track brain networks at a sub-second scale, providing important insights on the dynamics of those networks (Hassan and Wendling 2018; Kabbara et al. 2017). Since EEG is measured at the level of sensors distributed over the head, challenging computational problems need to be tackled to reconstruct the underlying networks with reasonable accuracy. In this context, the EEG source connectivity method enables identifying cortical networks while providing a satisfying trade-off between time and space resolutions (Hassan and Wendling 2018). However, many methodological questions related to the EEG source connectivity analysis remain unanswered. In fact, each step of the analysis involves different choices that may significantly affect the resulting functional network, which poses a key challenge to the topic of research reproducibility. Hence, the effect of several factors needs to be investigated and quantified for a more consistent and reliable use of this method.

First, several studies investigated the effect of the number of electrodes on EEG source localization in simulations in the context of epilepsy (Song et al. 2015; Sohrabpour et al. 2015; Goran Lantz et al. 2003). More specifically, it has been shown that the number of EEG electrodes has a direct influence on the localization error: a higher number of electrodes is associated with a significant decrease in localization error (Goran Lantz et al. 2003; Song et al. 2015; Sohrabpour et al. 2015). In both studies by (Goran Lantz et al. 2003) and (Sohrabpour et al. 2015), a dramatic decrease in localization error occured when increasing the number of electrodes from 32 to 64.

Another critical influencing factor in the EEG source connectivity pipeline is the algorithm chosen to solve the inverse problem, which is ill-posed due to its non-uniqueness and the instability of its solution (see (Grech et al. 2008) for a review). Several studies quantifying the performance of different inverse methods, in simulated and experimental EEG/MEG data, concluded that the choice of the inverse method significantly influences source estimation results (Anzolin et al. 2019; Mahjoory et al. 2017; Hedrich et al. 2017; Bradley et al. 2016; Grova et al. 2006; Halder et al. 2019; Tait et al. 2021; Allouch et al. 2022). However, no consistent conclusions have been made regarding one method that would stand apart from the others in terms of performance, which can also be related to the analyzed conditions.

The choice of the functional connectivity metric is also a critical step. A wide range of measures are used in the field, and each differs in the aspect of the data that is being investigated (amplitude-vs phase-based measures / directional vs non-directional connectivity, prone/robust to source leakage), (see (Friston 2011; Pereda, Quiroga, and Bhattacharya 2005; Cao et al. 2022) for a review), resulting in a significant variability of performance and interpretations (Colclough et al. 2016; H. E. Wang et al. 2014; Wendling et al. 2009; Hassan, Merlet, et al. 2017; Allouch et al. 2022).

In this context, simulation studies are of utmost interest, since they provide an otherwise inaccessible ground-truth for an objective evaluation of the methods/techniques under investigation. Several approaches have been suggested to provide such ground-truth. For instance, a toy model was used in (Schelter et al. 2006), where a signal was acting as an oscillator and was driving the activity of other structures. The use of Multivariate autoregressive (MVAR) models is also frequent (Anzolin et al. 2019; Haufe and Ewald 2016). However, such models are limited either in terms of their spectral properties, or in terms of their linearity and lack of complexity as compared to actual brain activity. As an attempt to bridge this gap, we used here a popular model for cortical dynamics, namely neural mass models (NMMs), to simulate resting state networks, and more specifically the default mode network (DMN) and dorsal attentional network (DAN). Then, we quantified the effect of three key factors involved in the EEG source connectivity analysis:

1. The EEG channel density (going from 19,32,64,128 to 256 channels).
2. The inverse method used to reconstruct EEG sources. We selected three of the widely used algorithms in the EEG/MEG community, namely i) weighted minimum norm estimate (wMNE) (Fuchs et al. 1999; Lin et al. 2006), ii) exact low-resolution brain electromagnetic tomography (eLORETA) (Pascual-Marqui 2007), and iii) linearly constrained minimum variance (LCMV) beamformer (Van Veen et al. 1997).
3. The functional connectivity metric. Since we did not intend to present an exhaustive comparison between all available metrics, we selected two phase- and two amplitude-based metrics: the phase-locking value (PLV) (Lachaux et al. 2000), phase-lag index (PLI) (Stam, Nolte, and Daffertshofer 2007) and amplitude envelope correlation (AEC) with and without source leakage correction (Hipp et al. 2012; Colclough et al. 2016, 2015). Finally, cortical networks computed for each combination of methods were quantitatively compared to the reference simulated networks for all experimental conditions investigated in this study.

## 2. Materials and Methods

A schematic diagram of the analysis pipeline is summarized in Figure 1.

**Figure 1.**
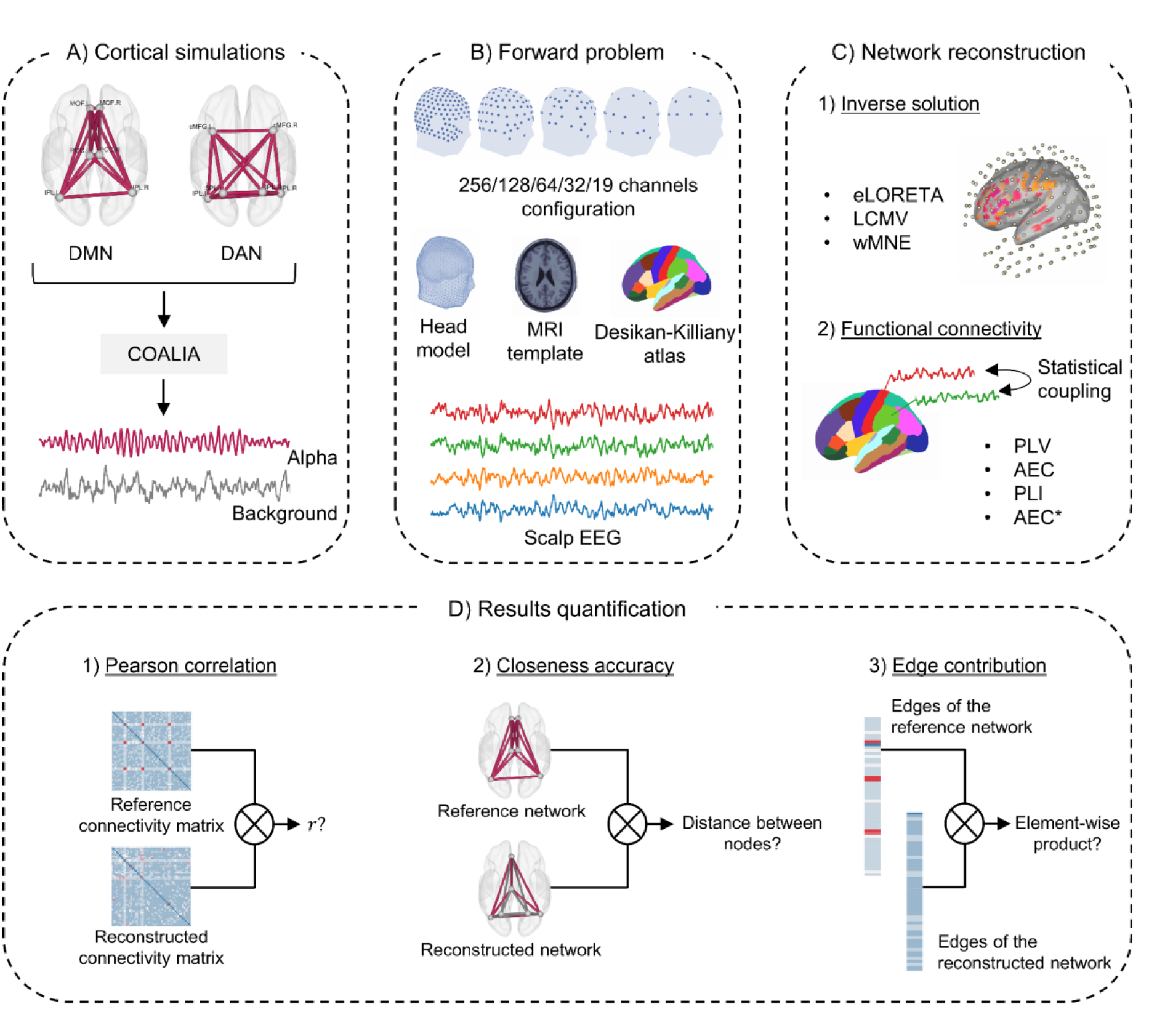
Pipeline used in the present study. (A) Generation of reference, ground-truth cortical-level electrophysiological signals using a realistic model of neuronal activity. (B) Projection of source-level activity on scalp-level EEG sensors by using an anatomically accurate head model. (C) Network reconstruction using different methods of source reconstruction based on scalp-level signals, and of statistical coupling between signals (functional connectivity metrics). (D) quantifying the level of matching between estimated and reference networks. *DMN - default mode network. DAN - Dorsal attentional network. eLORETA - exact low resolution electromagnetic tomography. LCMV - linearly constrained minimum norm beamforming. wMNE - weighted minimum norm estimate. PLV - phase-locking value. AEC - amplitude envelope correlation. PLI - phase-lag index. AEC* - amplitude envelope correlation with source leakage correction*.

### 2.1. Simulations

The simulated cortical networks (DMN and DAN) each included six regions based on the Desikan-Killiany atlas (Desikan et al. 2006) in terms of region parcellation. The DMN consisted of the right and left posterior cingulate cortex (PCC), medial orbitofrontal (MOF) gyrus, and inferior parietal lobe (IPL). Regarding the DAN, this network consisted of the right and left inferior parietal lobe (IPL), caudal middle frontal gyrus (cMFG), and superior parietal lobe (SPL). The choice of those regions was based on their frequent occurrence in previous resting-state studies (E. A. Allen et al. 2018; Elena A. Allen et al. 2014; Damoiseaux et al. 2006; M. D. Fox and Raichle 2007; Greicius et al. 2003; Baker et al. 2014; Shirer et al. 2012; Kabbara et al. 2017; Kabbara, Paban, and Hassan 2021). Cortical-level activity was generated using a flexible neural mass model framework, named COALIA. This multi-population neural mass model enables the simulation of brain-scale electrophysiological activity while accounting for the macro-(between regions) and micro-circuitry (within a single region) of the brain, with one neural mass representing the local field potential of one Desikan-Killiany atlas region [for details, readers may refer to (Bensaid et al. 2019)]. Activity in the alpha band ([8 − 12] *Hz*) was attributed to the regions belonging to reference RSNs, while background activity was assigned to remaining cortical regions. A variability between simulated data segments was introduced at the subject level, as well as at the level of epochs per subject. Each “virtual subject” had different connectivity matrices provided to the model, while each epoch for the same subject had a different input noise (*mean* = 90, *standard deviation* = 30) set within the model. More specifically, for each subject, a different fractional anisotropy matrix of the HCP dataset was used (Van Essen et al. 2013), and the weights corresponding to an RSN-connection were modified and set to a value of (1 ± 20%). A corresponding scaling of the matrices followed in accordance with COALIA’s requisites and the type of each input matrix (inhibitory/excitatory). A total of 50 “virtual subjects”, 4 epochs per subject (i.e., 200 data segments) were simulated; with a duration of 40 seconds each and a sampling rate of 2048 *Hz*. The time delay between NMMs was determined by the euclidean distance between the centroids of Desikan-Killainy’s regions divided by the velocity of action potentials propagation, which was set as 100 *cm/s*. An example of simulated cortical signals is shown in Supplementary Materials, Figure S1 (A).

### 2.2. Forward problem

Scalp EEG signals were estimated from simulated cortical activity by solving the forward problem as follows:

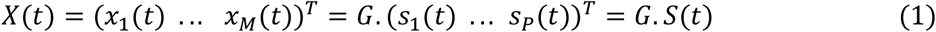

where *X*(*t*) represents scalp EEG signals, *S*(*t*) simulated cortical time series, and *G* the (*M* × *P*) Gain (leadfield) matrix. More specifically, *G* quantifies the contribution of each cortical source to the generation of scalp signals by taking into account the geometrical and electrical characteristics of the head. Here, the gain matrix was computed using a realistic head model (Colin27 MRI template) using the boundary element method (BEM), implemented within the OpenMEEG package (Gramfort et al. 2010) in the Fieldtrip toolbox (Oostenveld et al. 2011). The sensor space was defined based on the GSN HydroCel electrodes configuration (EGI, Electrical geodesic Inc) with 256, 128, 64, and 32 channels, as well as the international 10-20 system with 19 channels. The leadfield matrix used for solving the forward problem, as well as the inverse problem, was constrained to 66 leadfield vectors representing the contribution of the sources located at the centroid of the regions of interest defined by the Desikan-Killiany atlas (Desikan et al. 2006) (right and left insula were excluded, leaving 66 regions of interest). To mimic measurement noise, white gaussian noise was added to scalp EEG (Anzolin et al. 2019) as follows:

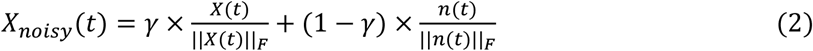

where *X*(*t*) and *n*(*t*) represent the scalp EEG signals and white uncorrelated noise signals, respectively; and ||. ||_*F*_ denotes their Frobenius norm. *γ* ranges from 0.1 to 1. An example of scalp EEG signals obtained for different *γ* values are shown in Supplementary Materials, Figure S1 (B). Results shown in the main manuscript correspond to *γ* = 1, i.e., no added measurement noise.

### 2.3. Inverse problem

The first step to reconstruct cortical networks was to estimate the dynamics of cortical sources from scalp EEG data, i.e., determining the position, orientation, and magnitude of dipolar sources *Ŝ*(*t*). Cortical sources were located at the centroids of Desikan-Killiany regions, and oriented normally to the cortical sheet. Thus, the inverse problem was reduced to computing the magnitude of dipolar sources *Ŝ*(*t*):

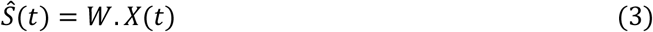

Several algorithms have been proposed to solve this problem, and estimate *W* based on different assumptions related to the spatiotemporal properties of sources and regularization constraints (see (Awan, Saleem, and Kiran 2019; Grech et al. 2008; Baillet, Mosher, and Leahy 2001) for a review). Inverse solutions can be classified into two major families: minimum norm estimates and beamformers. The former reconstructs all sources simultaneously by minimizing the difference between the data *X*(*t*) and predicted data *G*. *Ŝ*(*t*), while beamformers take an adaptive spatial-filtering approach in which each source is scanned independently. In this study, we focused on three commonly used solutions in EEG source reconstruction: weighted minimum norm estimate (wMNE) (Fuchs et al. 1999; Lin et al. 2006), exact low-resolution brain electromagnetic tomography (eLORETA) (Pascual-Marqui 2007), and linearly constrained minimum variance (LCMV) (Van Veen et al. 1997).

#### 2.3.1. Weighted minimum norm estimate (wMNE)

wMNE (Lin et al. 2006; Fuchs et al. 1999) is a derivative of the minimum norm estimate (MNE) (Hämäläinen and Ilmoniemi 1994), which proposes a solution that fits the measurements with a least square error. However, wMNE compensates further for the tendency of MNE to favor weak and surface sources:

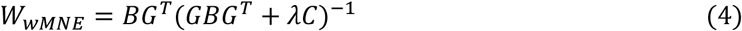

where *λ* is the regularization parameter, and *C* the noise covariance matrix (set to the identity matrix in our case). The matrix *B* is a diagonal matrix built from matrix *G* with non-zero terms inversely proportional to the norm of lead field vectors. This matrix adjusts the properties of the solution by reducing the bias inherent to the standard MNE solution:

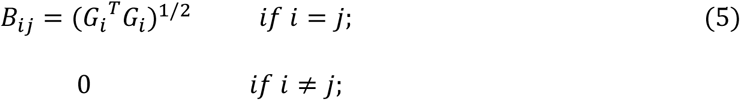

*R*egarding the regularization parameter *λ*, we used the recommended default value (1/SNR; SNR=3) included in the Brainstorm toolbox (Tadel et al. 2011).

#### 2.3.2. Exact low-resolution brain electromagnetic tomography (eLORETA)

eLORETA belongs to the family of weighted minimum norm inverse solutions. However, in addition to compensating for depth bias, it also has exact zero error localization in the presence of measurement and structured biological noise (Pascual-Marqui 2007):

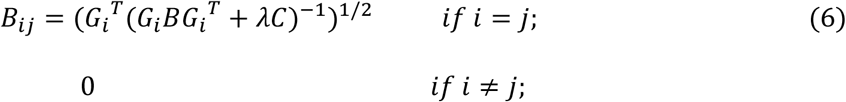

*R*egarding the regularization parameter *λ*, we used the default value (0.05) of the Fieldtrip toolbox (Oostenveld et al. 2011).

#### 2.3.3. Linearly constrained minimum-variance (LCMV) beamformer

The LCMV beamformer (Van Veen et al. 1997) takes an adaptive spatial-filtering approach and estimates the activity for a source at a given location while simultaneously suppressing contributions from all other sources and noise captured in the data covariance.

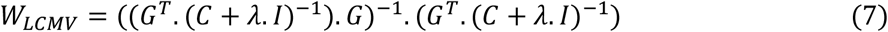

*T*he regularization parameter *λ* was set to 0.05.

### 2.4. Functional connectivity

Following the reconstruction of cortical dynamics, functional connectivity was assessed to estimate cortical networks. Two distinct approaches can be adopted to compute functional connectivity: phase-based and amplitude-based techniques. Here, we used the phase-locking value (PLV) and the phase-lag index (PLI) as an example of the former, and the amplitude envelope correlation (AEC) as an example of the latter.

#### 2.4.1. Phase-locking value (PLV)

For two signals *x*(*t*) and *y*(*t*), the phase-locking value (Lachaux et al. 2000) is defined as:

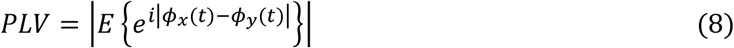

*w*here *E*{. } is the expected value operator, and *ϕ*(*t*) is the instantaneous phase derived from the Hilbert transform. PLV was computed over consecutive non-overlapping sliding windows, with the length of the window *δ* set to 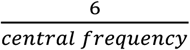 as recommended in (Lachaux et al. 2000), and where 6 is the number of cycles in a given frequency band. Thus, *δ* equals 600 *ms* in the considered alpha band ([8 − 12] *Hz*) where the central frequency was equal to 10 *Hz*. PLV values were then averaged over all sliding windows.

#### 2.4.2. Phase-lag index (PLI)

The PLI originally proposed in (Stam, Nolte, and Daffertshofer 2007) is a measure of the asymmetry of the distribution of phase differences between two signals. It aims at overcoming the issue of source leakage by discarding phase differences centered around 0 and *π*, that is, removing zero-lag connections. For two signals *x*(*t*) and *y*(*t*), PLI is defined as follows:

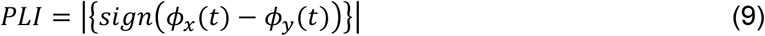

where *E*{. } is the expected value operator, and *ϕ*(*t*) is the instantaneous phase derived from the Hilbert transform. Similarly to PLV, PLI was computed over consecutive non-overlapping sliding windows (600 *ms*). PLI values were then averaged over all sliding windows.

#### 2.4.3. Amplitude envelope correlation (AEC)

AEC was computed as the Pearson correlation between the signals’ envelopes derived from Hilbert transform (Hipp et al. 2012; Brookes et al. 2011). Similar to PLV and PLI computation, a sliding window approach was adopted. Based on (O’Neill et al. 2017), the window length was set to 6 *s* with an overlap of 0.5 *s*.

#### 2.4.4. Amplitude envelope correlation with source leakage correction (AEC*)

Zero-lag signal overlaps caused by spatial leakage were removed using a symmetric orthogonalization approach detailed in (Colclough et al. 2015). Orthogonalization was applied over the entire data segments. Following the orthogonalization procedure, AEC was computed as previously described in section 2.4.3.

### 2.5. Results quantification and statistical testing

We used three metrics to assess the performance of different channel densities and the tested inverse solutions and connectivity measures.

#### 2.5.1. Pearson correlation

First, the Pearson correlation between the reference and reconstructed connectivity matrices was computed as a global measure of the similarity between networks. Matrices were not thresholded nor binarized.

#### 2.5.2. Closeness accuracy

Second, closeness accuracy (*CA*), defined as follows, was used for a node-wise analysis of the results:

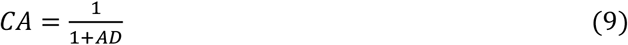

where *AD* is the average distance (in *cm*) between the reference and reconstructed networks given by:

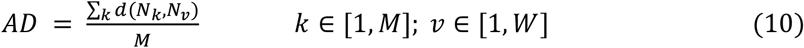

*w*here *d*(*N*_*k*_, *N*_*v*_) denotes the euclidean distance between the node *N*_*k*_ in the reconstructed network, and the nearest node *N*_*v*_ in the reference simulated network. *M* and *W* represent the total number of the nodes detected in the reconstructed and reference networks, respectively. Prior to computing the *CA*, networks were thresholded by keeping the edges with the highest 0.7% weight values, the choice of this proportion corresponding to the number of edges in the simulated RSNs (30 edges).

#### 2.5.3. Edge contribution

Finally, we investigated the contribution of individual edges to the correlation values obtained between the reference and reconstructed networks (Finn et al. 2015; Colclough et al. 2016). The set of edges of the reference and reconstructed (non-thresholded) networks were first z-score normalized (*mean* = 0, *std* = 1). Second, the edge contribution *φ* was calculated as the element-wise product between the two normalized edge vectors. Suppose that 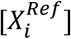 and 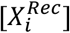 are the set of edges of the reference and reconstructed networks for an epoch *i*, after z-score normalization, then:

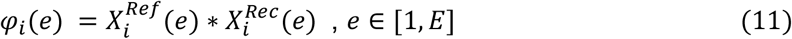

where *i* is the epoch index, *e* the edge index, and *E* the total number of edges, and the average of *φ*_*i*_ across all subjects and all epochs is denoted *ϕ*. High positive values in *ϕ* corresponds to edges that are consistent within- and between-subjects.

All statistical analyses were performed using R (R Core Team 2020). Linear mixed models implemented using the {*lme4}* package (Bates et al. 2015) were used to assess the effect of (i) number of sensors, (ii) inverse solution, and (iii) functional connectivity metric as fixed effects on correlation and closeness accuracy. The “subject” was added as a random intercept to account for simulation-related variability. Assumptions of normality and homoscedasticity of the model’s residuals were graphically checked. Regarding correlation, the following model was used:

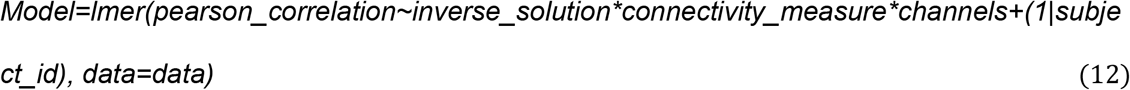

The same model was used for closeness accuracy with the difference that closeness accuracy was inverse-transformed because of a better compliance with the model’s assumptions in that case. Calculation of the significance of fixed effects was performed with F tests using the Anova function of the {*car*} package (J. Fox and Weisberg 2019), and post-hoc analyses were performed using z-tests with the *glht* function of the {*multcomp*} package (Hothorn, Bretz, and Westfall 2008) that provides corrected p-values. Marginal and conditional R^2^ were calculated with the {*MuMin*} package (Barton 2009) to estimate the variance explained by the models. The significance threshold was set at *p* = 0.05.

## 3. Results

Figure 2 and Figure 3 show, respectively, the distributions of correlation and closeness accuracy values, for all tested conditions (results obtained for DAN can be found in the Supplementary Materials, Figure S2, S3). Since the orthogonalization procedure is limited by the rank of the data (Colclough et al. 2015), it can only be applied in our case with 128 and 256 electrodes. Therefore, the statistical tests were repeated twice:

**Figure 2.**
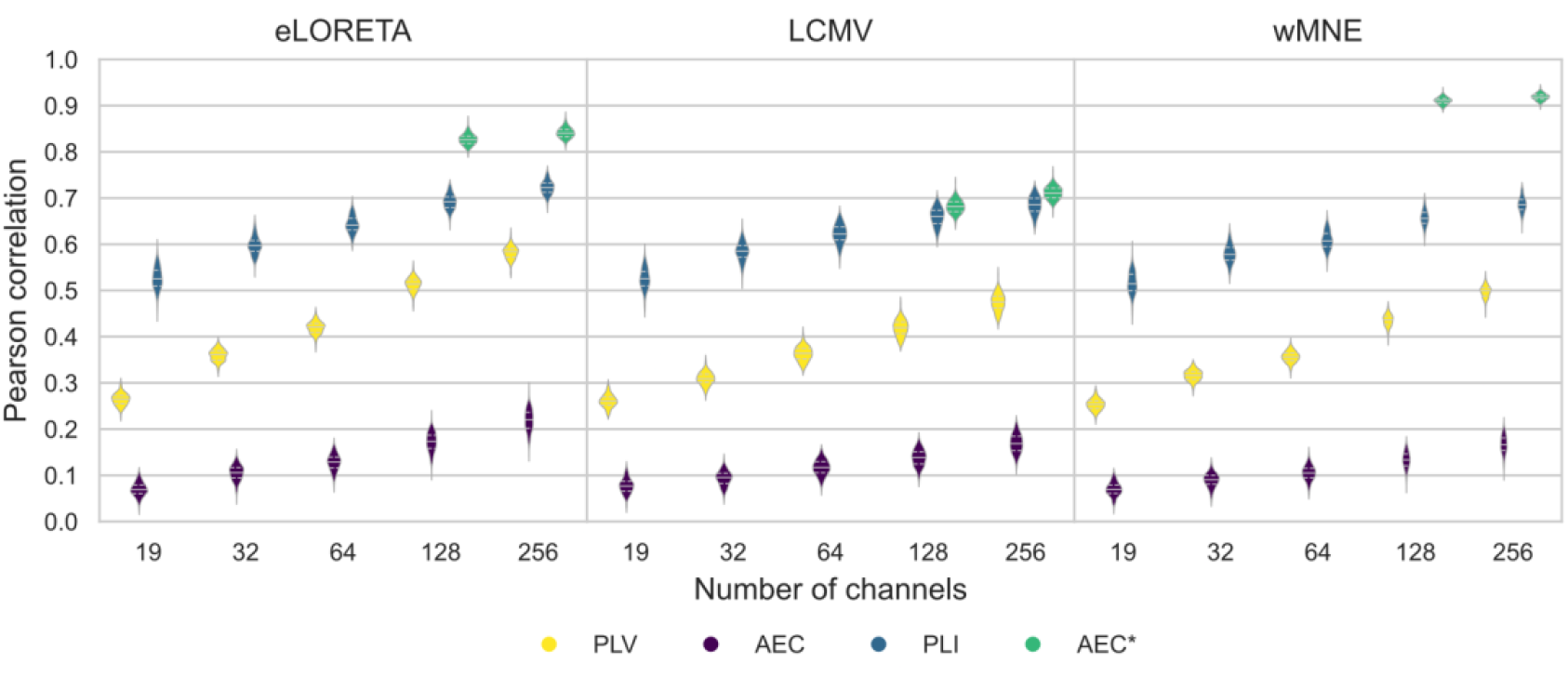
Violin plots of the Pearson correlation values computed between the reference and reconstructed DMNs for all electrode montages and inverse methods/connectivity metrics combinations. *eLORETA - exact low resolution electromagnetic tomography. LCMV - linearly constrained minimum norm beamforming. wMNE - weighted minimum norm estimate. PLV - phase-locking value. AEC - amplitude envelope correlation. PLI - phase-lag index. AEC* - amplitude envelope correlation with source leakage correction*.

**Figure 3.**
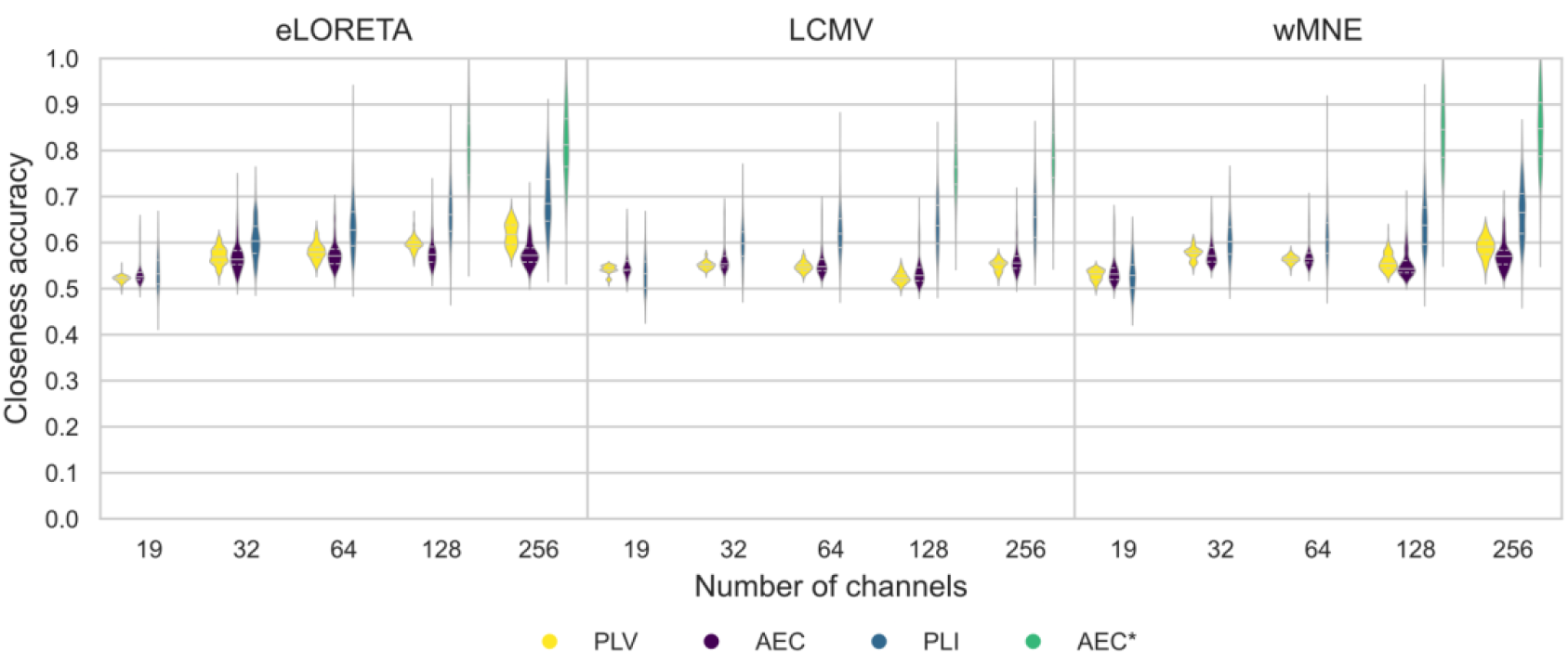
Violin plots of the closeness accuracy values computed between the reference and reconstructed DMNs for all electrode montages and inverse methods/connectivity metrics combinations. *eLORETA - exact low resolution electromagnetic tomography. LCMV - linearly constrained minimum norm beamforming. wMNE - weighted minimum norm estimate. PLV - phase-locking value. AEC - amplitude envelope correlation. PLI - phase-lag index. AEC* - amplitude envelope correlation with source leakage correction*.

- Case 1: comparing all electrodes configurations (19, 32, 64, 128, 256), inverse solutions (eLORETA, LCMV, wMNE), and three connectivity measures (PLV, AEC, PLI) where no orthogonalization is applied.
- Case 2: comparing two electrode configurations (128, 256), all inverse solutions (eLORETA, LCMV, wMNE) and all connectivity metrics (PLV, AEC, PLI, AEC*).

### 3.1. Pearson Correlation

For all inverse methods and connectivity metrics, the trend in correlation values was similar as the number of electrodes increased, for both DMN and DAN: a greater number of channels was associated with higher correlation values. Pearson correlation results demonstrate a drastic effect of increasing the number of electrodes on enhancing the accuracy of reconstructed networks, regardless of the chosen inverse solution/connectivity measure. Statistical tests comparing all channels configurations, inverse solutions and connectivity measures (PLV, AEC, PLI) (case 1) revealed that the effect of channel density was significant (F_(4, 8906)_ = 32256, p < 0.0001; full model marginal R^2^ = 0.99; conditional R^2^ = 0.99) as well as all pairwise comparisons (p < 0.0001). Similarly, comparing all inverse solutions and connectivity measures (PLV, AEC, PLI, AEC*) at 128 and 256 channels (case 2) revealed a significant effect of the number of electrodes (F_(1, 4727)_ = 6142, p < 0.0001; full model marginal R^2^ = 0.99; conditional R^2^ = 0.99). Significant effects of the inverse method (case 1: F_(2, 8906)_ = 4388, p < 0.0001; case 2: F_(2,4727)_ = 10396, p < 0.0001), functional connectivity measure (case 1: F_(2, 8906)_ = 651672, p < 0.0001; case 2: F_(3, 4727)_ = 371598, p < 0.0001) were also identified. Moreover, reconstructed networks differed as a function of the inverse solution/connectivity metric combination (case 1: F_(4, 8906)_ = 548, p < 0.0001; case 2: F_(6, 4727)_ = 4858, p < 0.0001). *Post-hoc* analysis showed significant differences between inverse solution algorithms (case 1: p<0.001, except for eLORETA *vs* wMNE: p=0.17; case 2: p<0.001 except for LCMV *vs* wMNE: p = 0.01), as well as connectivity measures (p<0.001). Pairwise comparisons of the different combinations of inverse solution and functional connectivity methods also revealed significant differences between combinations (case 1: p<0.001 except for wMNE/AEC vs eLORETA/AEC (p = 1), LCMV/PLI *vs* eLORETA/PLI (p = 0.99) and LCMV/PLV *vs* eLORETA/PLV (p = 0.48); case 2: p<0.001 except for wMNE/AEC *vs* LCMV/AEC (p = 0.13) and wMNE/PLI *vs* LCMV/PLI (p = 0.97). At high sensor densities, AEC computed between orthogonalized time series along with wMNE and eLORETA, exhibited the best reconstruction accuracy. PLI, which also discards zero-lag connections, came second with all three inverse solutions and regardless of channel density. Regarding measures which do not compensate for spatial leakage, PLV had a moderate performance, whereas correlation values obtained with AEC were systematically low, regardless of the inverse method and number of electrodes.

### 3.2. Closeness accuracy

Differences in closeness accuracy values between evaluated electrode configurations were not as clear as with the Pearson correlation values, except for PLI and AEC*, which were associated with a higher between-epochs variability. Similar to correlation results, closeness accuracy values were significantly affected by the number of electrodes (case 1: F_(4, 8906)_ = 1278, p < 0.001; full model marginal R^2^ = 0.43; conditional R^2^ = 0.44; case 2: F_(1, 4727)_ = 267, p < 0.001; full model marginal R^2^ = 0.81; conditional R^2^ = 0.82), inverse method (case 1: F_(2, 8906)_ = 304, p < 0.001; case 2: F_(2, 4727)_ = 395, p < 0.001), connectivity metric (case 1: F_(2, 8906)_ = 2167, p < 0.001; case 2: F_(3, 4727)_ = 6696, p < 0.001), and inverse method/connectivity metric combination (case 1: F_(4, 8906)_ = 53, p < 0.001; case 2: F_(6, 4727)_ = 74, p < 0.001). Pairwise comparisons of channel density showed significant differences between results obtained with 19 and (32, 64, 128, 256) channels (p<0.001), as well as between 32 and 128 electrodes (p = 0.01). Other pairwise comparisons of electrode configurations were not significant. When comparing all electrodes configurations, inverse solutions and (PLV, AEC, PLI), *post-hoc* analysis showed significant differences between inverse solutions (p<0.001) except for wMNE *vs* eLORETA (p = -1.79), as well as between connectivity metrics (p<0.001) except for PLI *vs* AEC (p = 1). Regarding the inverse solution/connectivity measure combination, differences were not all significant. When comparing results obtained with 128 and 256 electrodes and all inverse solutions and connectivity measures, significant differences were obtained for all inverse solutions (p<0.001), connectivity measures (p<0.001) and inverse/ solution connectivity measure combination (p<0.001 except for wMNE/PLV *vs* WMNE/AEC (p = 0.89) and wMNE/PLI *vs* LCMV/PLI (p = 1). Detailed statistical tests results relative to the DMN and DAN are reported in the Supplementary Materials.

### 3.3. Edge contribution

In Figure 4 and Figure 5, the contribution of DMN edges averaged across all subjects and epochs are presented for all sensor densities and inverse method/connectivity metric combination (results for the DAN are shown in Figure S4 and S5 in Supplementary Materials). The contribution of DMN connections to the correlation value computed between the reference and reconstructed networks increased when increasing the number of electrodes. The averaged edge contribution measure can be seen as a reflection of the consistency of DMN connections in the reconstructed networks across all data segments, i.e., a higher number of electrodes is associated with a greater consistency of DMN edges. This trend was clearer with phase-based connectivity measures (PLV and PLI) than with AEC.

**Figure 4.**
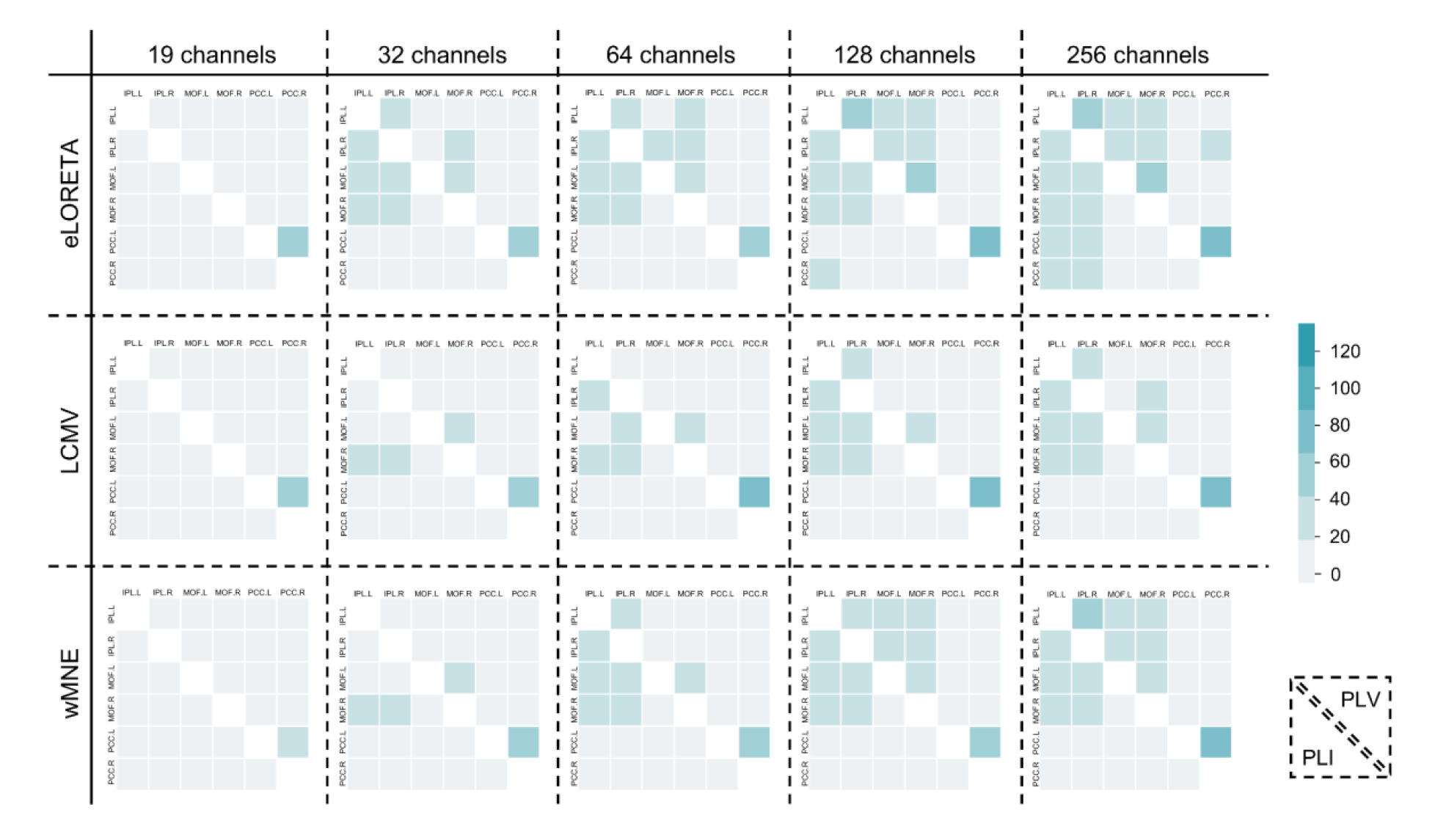
Heatmaps of contribution of the DMN edges (see Materials and Methods section) averaged across all subjects and epochs are shown for all sensor densities and inverse method/connectivity measure (PLV, PLI) combinations. *eLORETA - exact low resolution electromagnetic tomography. LCMV -linearly constrained minimum norm beamforming. wMNE - weighted minimum norm estimate. PLV - phase-locking value. AEC - amplitude envelope correlation. PLI - phase-lag index. AEC* - amplitude envelope correlation with source leakage correction*.

**Figure 5.**
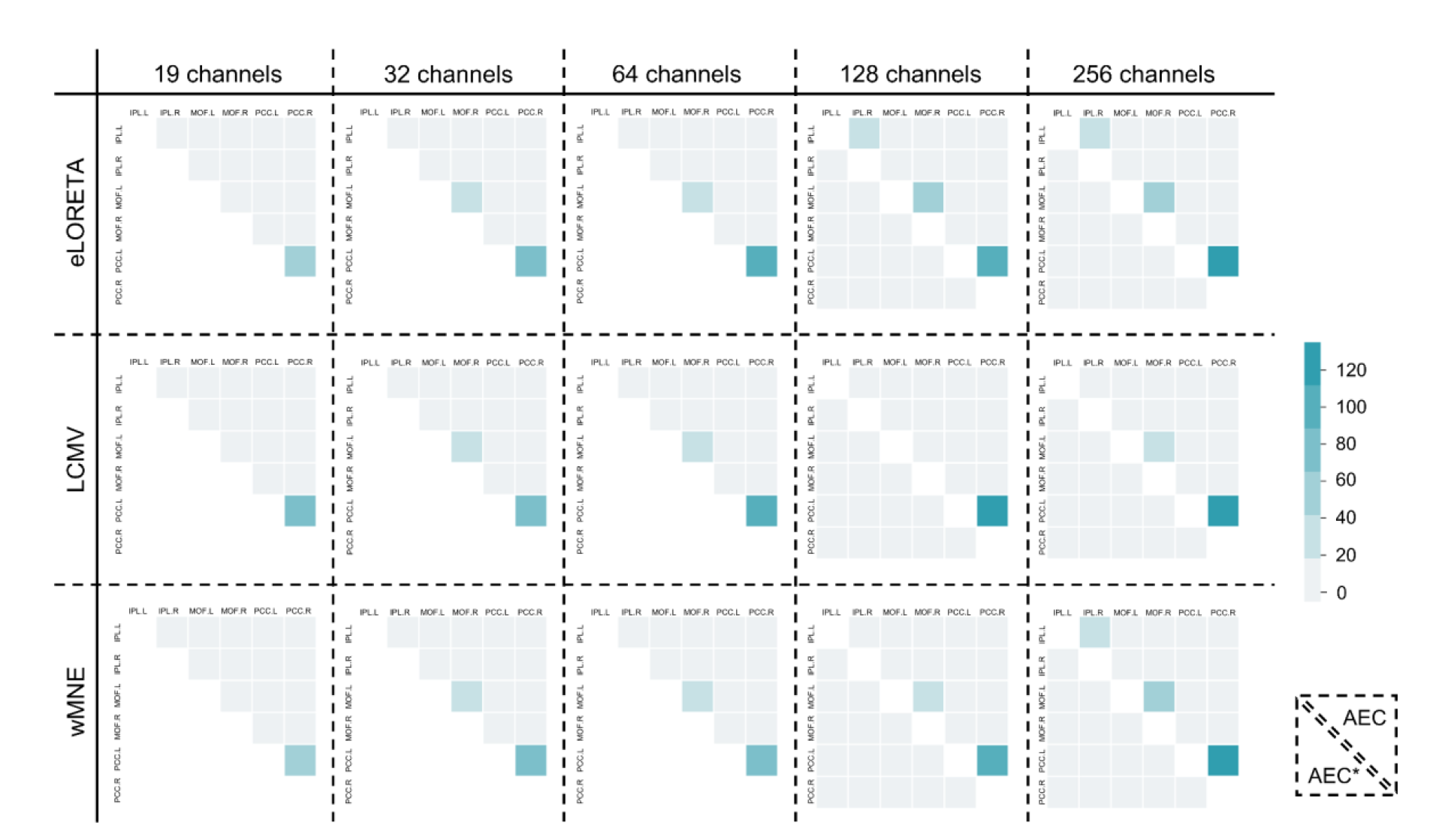
Heatmaps of contribution of the DMN edges averaged across all subjects and epochs are shown for all sensor densities and inverse method/connectivity measure (AEC, AEC*) combinations. *eLORETA - exact low resolution electromagnetic tomography. LCMV - linearly constrained minimum norm beamforming. wMNE - weighted minimum norm estimate. PLV - phase-locking value. AEC - amplitude envelope correlation. PLI - phase-lag index. AEC* - amplitude envelope correlation with source leakage correction*.

We then thresholded the averaged edge contribution matrices and kept only the 30 edges (number of edges in the reference RSNs) having the highest contribution values. The percentile of edges belonging to DMN (DAN) are thus shown in Figure 6 (Figure S6 in Supplementary Materials). With PLV and AEC, the percentage of edges within DMN increased at higher channel density with AEC having percentage values not exceeding 50%. In contrast, the majority of edges with the highest contribution were within the DMN when using PLI or AEC with source leakage correction regardless of the number of channels.

**Figure 6.**
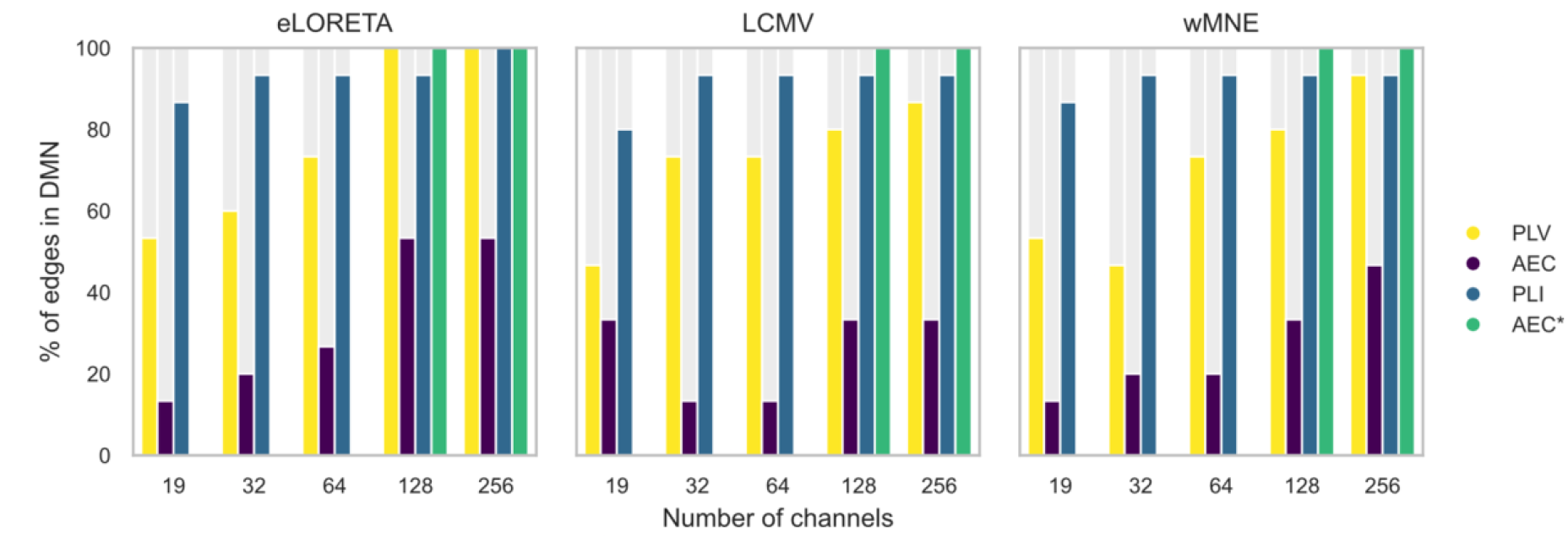
Barplots of the percentage of edges located within the DMN among the edges having the ≈ 0.7% highest contribution values. *eLORETA - exact low resolution electromagnetic tomography. LCMV linearly constrained minimum norm beamforming. wMNE - weighted minimum norm estimate. PLV - phase-locking value. AEC - amplitude envelope correlation. PLI - phase-lag index. AEC* - amplitude envelope correlation with source leakage correction*.

### 3.4. Effect of white measurement noise

In Figure 7, we plotted the Pearson correlation (mean+standard deviation) as a function of the measurement noise level (number of channels = 256). eLORETA and wMNE had similar performances. They both exhibited stable correlation values (≈ 0.7) for *γ* ≥ 0.4 with PLV and *γ* ≥ 0.6 with PLI. In contrast, LCMV combined with PLV and PLI had a linear trend where correlation values increased when added measurement noise decreased. Interestingly, AEC* had the highest performance in the absence of noise (*γ* = 1). However, its performance degrades drastically when noise is added. eLORETA/AEC, wMNE/AEC and LCMV/AEC also were stable in terms of performance, albeit performance being low. (Results corresponding for DAN are shown in Figure S7 in Supplementary Materials).

**Figure 7.**
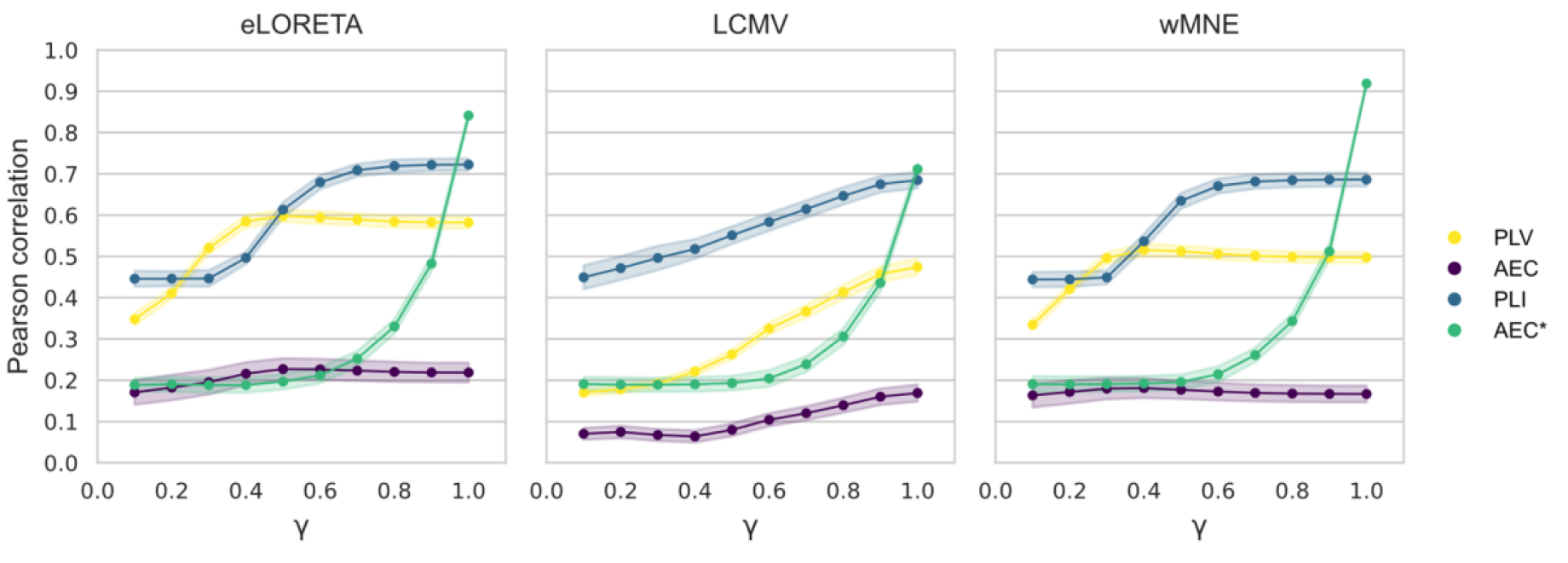
Mean and standard deviation of the Pearson correlation computed between the reference and reconstructed DMNs for different levels of measurement noise using 256 channels. *eLORETA - exact low resolution electromagnetic tomography. LCMV - linearly constrained minimum norm beamforming. wMNE - weighted minimum norm estimate. PLV - phase-locking value. AEC - amplitude envelope correlation. PLI - phase-lag index. AEC* - amplitude envelope correlation with source leakage correction*.

## 4. Discussion

The EEG/MEG source connectivity technique has gained increased interest due to its ability to reconstruct functional networks in the cortical space with a high temporal resolution. As aforementioned, however, there is still no agreement to date over a unified detailed EEG source connectivity pipeline, and several parameters/methods are involved which require cautious tuning. Here, our objective was to evaluate some of the parameters related to the EEG source connectivity analysis in the context of resting state networks, and investigate the variability in the results caused by different choices in the analysis pipeline. We mainly focused on the effect of the number of electrodes, inverse method, and functional connectivity metric. In a recent study, we tested the effect of the number of electrodes, and tested wMNE and eLORETA along with PLV and wPLI in the context of simulated epileptiform activity (Allouch et al. 2022). In this specific case, epileptiform signals had a sufficiently high signal-to-noise-ratio (SNR) to be distinguished from background noise (Wa 1983; Iwasaki et al. 2005), which might have facilitated the identification of underlying networks. However, a growing interest in the last two decades has led to an increased number of resting-state studies where the significantly lower SNR could be in favor of different methods/parameter tuning in EEG source connectivity analysis. Therefore, in the present study, we used neural mass models to simulate resting-state brain activity (DMN and DAN) in the cortical space, and derived the corresponding scalp EEG signals by solving the forward problem. Then, we reconstructed the corresponding cortical networks. This pipeline was repeated using different electrode densities (19, 32, 64, 128, 256 channels) to solve the forward problem, three algorithms to solve the inverse problem (wMNE, eLORETA, LCMV), and four metrics to assess functional connectivity (PLV, AEC, PLI, leakage-corrected AEC (AEC*)), at different levels of measurement noise.

### 4.1. Number of electrodes

To the best of our knowledge, the effect of the number of electrodes on the reconstruction of EEG-based resting-state cortical networks has never been studied before, especially in the presence of a ground-truth enabling an objective comparison of the tested electrode configurations. Our results demonstrate a key role of the number of electrodes on the EEG source connectivity analysis: a more accurate reconstruction of cortical activity was achieved using high-density EEG (hd-EEG). This result is in line with previous simulations and empirical evidence (Song et al. 2015; Sohrabpour et al. 2015; Goran Lantz et al. 2003), as well as with the theoretical foundation (Srinivasan, Tucker, and Murias 1998; Song et al. 2015). As established in (Srinivasan, Tucker, and Murias 1998), an accurate characterization of the spatial electrophysiological information requires a higher number of electrodes. A high inter-electrode distance (i.e., corresponding to a low number of EEG electrodes) can induce aliasing, and therefore high spatial frequency signals are misrepresented as low spatial frequency signals due to the violation of the Nyquist criteria (*F*_*s*_ > 2 × *F*_*max*_) (Srinivasan, Tucker, and Murias 1998; Song et al. 2015). In (Song et al. 2015) results were found to be independent of the inverse method (minimum norm/standardized low resolution brain electromagnetic tomography). On the other hand, (Goran Lantz et al. 2003) showed that 63 electrodes were sufficient for a decent source localization with EPIFOCUS (a linear inverse solution that optimally localize single focal sources (Grave de Peralta Menendez et al. 2001; G. Lantz et al. 2001)), however, 100 electrodes were required when using the weighted minimum norm estimate. Consistently, (Goran Lantz et al. 2003) and (Sohrabpour et al. 2015) showed that a dramatic decrease in localization error was achieved when increasing the number of electrodes from 32 to 64 electrodes. Based on our results, we suggest a minimum of 64 electrodes to be used in the course of EEG source reconstruction in the specific context of resting-state networks.

### 4.2. Inverse solution and connectivity measures

According to the source connectivity estimation, a plethora of methods offers the possibility to 1) reconstruct the dynamics of cortical activity, and 2) assess the functional connectivity between reconstructed sources. However, there is no consensus over an optimal approach, nor on whether such “best” technique exists. Both in terms of the correlation between reference and reconstructed networks, and in terms of the closeness accuracy of nodes detected in the reconstructed networks, our results showed that eLORETA and wMNE performed significantly better than LCMV, and that compensating for spatial leakage (PLI, AEC*) resulted in increased accuracy, regardless of the inverse solution. Significant discrepancies were observed with the different combinations of inverse methods and connectivity measures tested in this study, which was also observed in other comparative studies. In general, there is a lack of consistency across studies comparing several inverse methods and connectivity measures. For example, (Anzolin et al. 2019) showed that LCMV had a better performance globally as compared to eLORETA. Similarly, in (Mahjoory et al. 2017), a relatively strong difference was found between LCMV beamformer on one hand, and eLORETA/wMNE solutions on the other hand. In (Hedrich et al. 2017) the coherent maximum entropy on the mean (cMEM) showed similar localization error to MNE, dynamic statistical parametric mapping (DSPM), sLORETA, but lower spatial spread and reduced crosstalk. In (Bradley et al. 2016), the use of LORETA for source localization outperformed sLORETA and minimum norm least square (MNLS). Following an extensive comparison between six inverse methods, (Grova et al. 2006) recommended taking into account results from different methods when localizing actual interictal spikes. Results of source localization in (Halder et al. 2019) did not identify a clear winner between LCMV, eLORETA, MNE, DISC. (Tait et al. 2021) summarized the conditions where each method can be recommended, following a comparison of six inverse methods in resting state MEG data. It is noteworthy to mention that the discrepancy of the results was mainly related to the mathematical and physical constraints imposed by each of the inverse solutions. More precisely, wMNE searches for a solution with minimum power (Hämäläinen and Ilmoniemi 1994), while eLORETA tends to achieve exact zero error localization in the presence of measurement and structured biological noise (Pascual-Marqui 2007). On the other hand, LCMV beamformer assumes that only one dipole is active at a time, and tries to minimize the corresponding output energy (Van Veen et al. 1997).

In terms of functional connectivity metrics, (Colclough et al. 2016) assessed the consistency of different measures in experimental MEG resting state data and recommended using the correlation between orthogonalised, band-limited, power envelopes (AEC). On the other hand, following extensive simulation studies, (H. E. Wang et al. 2014) and (Wendling et al. 2009) both concluded that there is no ideal “one-fits-all” method for all data types: it is rather suggested to evaluate which conditions are appropriate for each method. In (Hassan et al. 2014) and (Hassan, Merlet, et al. 2017) in the context of epilepstic spikes, wMNE combined with PLV had better accuracy as compared to other algorithms. Let us note that, in (Allouch et al. 2022), wMNE combined with wPLI performed better in the context of epileptiform activity simulations.

A key limitation of those numerous studies (and ours) is that different methods were tested in different contexts and using different data types, which complicates further the identification of clear and concise guidelines on the topic. Thus, we are aware that, even with our additional contribution, we are still far from a generalization of the results obtained in the specific context tested in this study (simulated signals, resting state activity, alpha rhythms, number/location of cortical sources, etc…). In fact, it is entirely possible (and even probable) that the inverse solution/connectivity measure combination is context-specific, and that no ideal method can account for all data types or all research questions. However, this raises the question of whether it is possible to determine the method that is the most adapted in each context, which requests extensive investigation far beyond simple comparative studies. Meanwhile, cross-validation of the results using several methods/measures could be a reasonable compromise.

### 4.3. Methodological considerations

Taking together the conclusions from the vast majority of analysis and modeling choices, the identified networks might be also sensitive to other factors that were not investigated in this study. For instance, spatial resolution in the cortical space (i.e., number/size of parcellated ROIs) could affect the accuracy of reconstructed networks. Our simulations were all restricted to 66 ROIs (Desikan Killiany atlas) due to the model design. Many graph-based studies have reported that different parcellation scales resulted in significant differences in network parameters (clustering coefficient, characteristic path length, local and global efficiency, degree distribution, etc…), while inferences about small-world and scale-free properties were maintained across different scales (Hayasaka and Laurienti 2010; J. Wang et al. 2009; Zalesky et al. 2010; Fornito, Zalesky, and Bullmore 2010). In this study, we did not investigate how spatial resolution in the source space (i.e., number/size of ROIs) could affect the accuracy of reconstructed networks. Therefore, it would be interesting to evaluate if the optimal number of ROIs depends on the number of recording electrodes and *vice versa*, i.e., whether there exists an optimal ratio (number of electrodes/number of ROIs) that exhibits higher accuracy regarding source connectivity. Moreover, the effect of parcellation is not limited to the number of regions, but also involves the algorithms by which sources belonging to the same ROI are aggregated (averaging signals across all vertices, averaging their absolute values, power signals, keeping the first mode of PCA decomposition, choosing the maximum value among vertices at each time point…).

Here, we simulated two widely studied RSNs (DMN, DAN). The major difference between those two networks is the position of simulated sources (i.e., network’s regions). While the DMN exhibits a less distributed architecture with small to moderate distance between regions (especially the right and left MPFC and PCC), DAN regions are widespread across the cortex with higher inter-region distances. Due to this difference, we were able to test whether the results are affected by the specific spatial location of sources. However, the similarity of results obtained with both networks confirm, to some extent, the absence of a bias caused by the position of sources. For simplicity, we restricted the number of DMN and DAN regions in our simulations to six, and focused on the most consistent regions reported in the literature. However, other regions could be involved in the simulated DMN, such as the precuneus, isthmus cingulate, rostral anterior cingulate and lateral orbitofrontal cortex. The DAN could also include the middle temporal gyrus and frontal regions, such as the parsopercularis, parsorbitalis and parstriangularis (Kabbara et al. 2017; Kabbara, Paban, and Hassan 2021). An interesting future prospect would be testing the consistency of results when more complex networks are involved.

In the simulations used for this study, alpha activity [8-12] *Hz* was assigned to RSN, while background activity was attributed to the remaining regions. We acknowledge that such contrast between those two types of activity is not ideally realistic. Despite a potential significant interest, we were not able to simulate brain-like broadband signals covering all frequency bands, due to intrinsic model limitations. In addition to broadband simulations, frequency bands other than alpha could be tested with the aim to replicate those results. Another limitation of our study is our “static” approach for brain networks identification (i.e., one network estimated per epoch), which we could improve upon by investigating the accuracy of brain networks dynamics. Importantly, understanding the dynamics of brain networks provides crucial insights about brain functions both in resting-state (Hipp et al. 2012; F. de Pasquale et al. 2018; De Pasquale et al. 2016; E. A. Allen et al. 2018; Elena A. Allen et al. 2014; Baker et al. 2014; Kabbara et al. 2017; Kabbara, Paban, and Hassan 2021) and task-related paradigms (Shine et al. 2016; O’Neill et al. 2017; Elton and Gao 2015; Braun et al. 2015; Krienen, Yeo, and Buckner 2014; Fong et al. 2019; Hassan et al. 2015).

As aforementioned, inverse solutions and connectivity algorithms require tuning several parameters, such as the regularization parameter in wMNE, eLORETA, LCMV and the time window over which connectivity measures are computed. Here, we applied either default or usually used values proposed by the Fieldtrip and Brainstorm toolboxes, or values proposed in previous studies. To test the stability of reconstructed networks across different regularization values, we computed the Pearson correlation between the reference and reconstructed networks for different regularization values (PLV, 256 channels) and presented the results in Figure S8 in the Supplementary Materials. A decrease in the correlation values was observed when increasing the regularization of the inverse solution, emphasizing the importance of tuning the regularization parameter rather than using default values. Several methods are available to address this issue, falling into two general categories: those based on the estimation of the measurement noise, and those which are not (L-curve method, general-cross validation method, composite residual and smoothing operator, minimal product method, zero crossing) (Grech et al. 2008). An additional crucial factor that may introduce variability between results, and that emerges when dealing with experimental EEG data, is the pre-processing procedure that precedes network reconstruction. EEG signals are indeed usually contaminated by several artifacts and noise sources, that may each introduce undesired changes in the measurements and affect the signal of interest (Urigüen and Garcia-Zapirain 2015), especially in clinical, pediatric and aging populations (Pedroni, Bahreini, and Langer 2019). In order to clean EEG signals, pre-processing algorithms usually impose different constraints for accepting and rejecting artifactual epochs, and propose different techniques to eliminate artifacts. Hence, an important methodological consideration would be to investigate and systematically quantify the variability induced by different data preprocessing techniques.

The choice of the most convenient metric for quantifying the results is also a challenge. Here, in order to quantify the performance of the tested parameters, we used the (1) Pearson correlation computed between the reference and reconstructed networks, (2) closeness accuracy reflecting the distance between the nodes detected in the reconstructed network and those in the reference network, and, (3) averaged contribution of network edges to the correlation between connectivity matrices (i.e.,the consistency of DMN/DAN connections across all simulation epochs) and the percentage of edges having the highest contributions and falling within the simulated RSN. It should be noted that each metric measures different aspects of data, and it is important in the context of comparative studies to address the relevant aspect of data to prevent biases. In our case, the general trend in the results was preserved across all used metrics. While both Pearson correlation and closeness accuracy showed significant differences between tested parameters, the latter exhibited higher variability between epochs. Other metrics such as networks-based metrics have also been developed to evaluate the distance/similarity between networks, and a promising approach would be testing whether the differences between networks are maintained across different aspects investigated by such metrics (global analysis, edge-wise, node-wise, spectral graph analysis, Edit distance, kernel methods) (Mheich, Wendling, and Hassan 2020).

Importantly, correlation results presented in this article were computed between unthresholded weighted connectivity matrices. However, selecting a set of network edges for subsequent analysis while discarding others remains a debatable subject in the network neuroscience community. We usually face two main questions: 1) what method to use to perform thresholding, and 2) the choice of the threshold value when needed. Since we aimed to reduce further data manipulation and subjective intervention in the analysis, no threshold was applied prior to the computation of Pearson correlation and edge contribution matrices in our study. However, we also tested the effect of a proportional threshold (1%) on correlation values and reported the results in Figure S9 in Supplementary materials. Although lower correlation values and higher between-epochs variability were obtained, the general trend was similar to that of unthresholded matrices.

## 5. Conclusion

To sum-up, we simulated EEG data corresponding to RSNs and tested the effect of several key parameters (number of EEG electrodes and inverse solution/connectivity measure combination) on network reconstruction accuracy. Different analytical choices led to a high variability in the resulting networks. In the context of RSN simulations, our results demonstrate, as expected, that an accurate cortical network reconstruction requires a high number of EEG electrodes. Therefore, we recommend using hd-EEG (>64 channels) to infer cortical dynamics from recorded scalp EEG signals. In addition, we suggest a very careful choice of the inverse solution/connectivity measure combination, since our results highlight a significant variability in the networks reconstructed using different inverse solutions and connectivity measures. Such methodological variability and absence of analyses standardization represent a critical issue for neuroimaging studies that should be prioritized.

## Data and codes availability

Data simulated for this study are available at https://doi.org/10.5281/zenodo.6597385. Codes supporting the results of this study are available at https://github.com/sahar-allouch/var-RSNs. We used Matlab (Matlab 2018), Brainstorm toolbox (Tadel et al. 2011), Fieldtrip toolbox ((Oostenveld et al. 2011); http://fieldtriptoolbox.org), OpenMEEG (Gramfort et al. 2010) implemented in fieldtrip, R (R Core Team 2020) for statistical analysis, and Seaborn (Waskom 2021) and Matplotlib (Hunter 2007) for visualization.

## Acknowledgments

The authors would like to acknowledge the Lebanese National Council for Scientific Research (CNRS-L), the Agence Universitaire de la Francophonie (AUF) and the Lebanese University for granting Ms. Allouch a doctoral scholarship. This work is also supported by the Labex Cominlabs project PKSTIM. This work was also supported by the Institute of Clinical Neuroscience of Rennes (Projects named EEGCog and EEGNET3). Authors would like to thank Campus France, Programme Hubert Curien CEDRE (PROJET N° 42257YA) and the Lebanese Association for Scientific Research (LASER) for their support.

