## Supplementary Materials for "Effect of channel density, inverse solutions and connectivity measures on EEG resting-state networks: a simulation study"

### COALIA: a physiologically-inspired computational model

COALIA is a recently developed physiologically-grounded computational model (Bensaid et al. 2019) of large-scale brain activity. Using a bottom-up approach taking into account the detailed circuitry between the main neuronal subtypes, and anatomical regions from a widely used atlas, COALIA generates brain-scale electrophysiological activity while accounting for the macro- as well as the micro-circuitry of the brain. The basic unit of the model is the neural mass, a local network involving different neuronal types in which the electrical activity is averaged over the cells of a similar type, instead of describing individual cell dynamics as in microscopic models. Therefore, the neural mass model (NMM) is a mesoscopic model describing synchronized activity in local networks in which the micro-circuitry can be taken into account. At the level of a single neural mass, the model includes glutamatergic pyramidal neurons and three different types of GABAergic interneurons with physiologically-based kinetics (fast vs. slow). At the brain-scale level, each neural mass represents the local field activity of one region of the Desikan-Killiany atlas (66 regions; right and left insula were excluded) (Desikan et al. 2006). Given that each neural mass simulates the activity of one specific brain region, neural masses are then synaptically connected through long-range glutamatergic projections. This neuro-inspired model can simulate both cortical and thalamic activity. In the following, a brief description of the local NMM of neocortical is presented.

The neocortical module involved pyramidal cells (PCs) and three types of inhibitory GABAergic interneurons, namely, (1) somatic-targeting parvalbumin positive (PV+) basket cells (BC); (2) the dendritic-targeting somatostatin positive (SST) interneurons; and (3) vasoactive intestinal-peptide (VIP) expressing interneurons. BC and SST received excitatory inputs from PCs that are reciprocally inhibited by both of them. Pyramidal collateral excitation was implemented *via* an excitatory feedback loop passed by a supplementary excitatory population (PC') analogous to PC, except that it projected only to the subpopulation PC and received projection from PC as well. An inhibitory feedback loop was implemented to account for direct PV+/PV+ coupling through electrical gap-junctions. Communication through disinhibition was modeled by inhibitory projections first from VIP to SST, then from SST to BC. The non-specific influence from neighboring and distant populations was modeled by a Gaussian input noise corresponding to an excitatory input $p_c^n(t)$ that globally described the average density of afferent action potentials.

At the brain-scale, glutamatergic PCs originating from a single cortical column targeted PCs of other cortical columns by common feedforward excitation and GABAergic cells by disynaptic cortico-cortical feedforward inhibition. Variable time delays between NMMs were introduced in order to account for activity propagation delays caused by long-range connections.

**Table S1** COALIA simulation parameters (values and interpretation)

| ***Cortical module*** | | |
|---|---|---|
| **Parameter** | **Value** | **Interpretation** |
| $A_c$ | 4 mV | Amplitude of the cortical average EPSP |
| $B_c$ | 30 mV for background activity<br>9 mV for alpha rhythms | Amplitude of the cortical average IPSP ($GABA_{A,\ slow}$ mediated currents) |
| $G_c$ | 50 mV | Amplitude of the cortical average IPSP ($GABA_{A,\ fast}$ mediated currents) |
| $D_c$ | 100 mV | Amplitude of the cortical average IPSP ($GABA_{A,\ slow}$ mediated currents) |
| $\mathbf{1}/a_c$ | 1/100 s | Time constant of cortical glutamate-mediated synaptic transmission |
| $\mathbf{1}/b_c$ | 1/30 s | Time constant of cortical GABA-mediated synaptic transmission ($GABA_{A,\ slow}$ receptors) |
| $\mathbf{1}/g_c$ | 1/150 s | Time constant of cortical GABA-mediated synaptic transmission ($GABA_{A,\ fast}$ receptors) |
| $\mathbf{1}/d_c$ | 1/20 s | Time constant of cortical GABA-mediated synaptic transmission ($GABA_{A,\ slow}$ receptors) |
| $\mu_c$, $\sigma_c$ | 90 $s^{-1}$, 30 $s^{-1}$ | Mean and standard deviation of nonspecific cortical input |
| $C^n_{P,P'}$ | 135 | Collateral excitation connectivity constant of $n^{th}$ cortical population |
| $C^n_{P',P}$ | 100 | Collateral excitation connectivity constant of $n^{th}$ cortical population |
| $C^n_{BC,P}$ | 50 | BC to PC connectivity constant of $n^{th}$ cortical population |
| $C^n_{SST,P}$ | 20 | SST to PC connectivity constant of $n^{th}$ cortical population |
| $C^n_{P,BC}$ | 50 | PC to BC connectivity constant of $n^{th}$ cortical population |
| $C^n_{P,SST}$ | 50 | PC to SST connectivity constant of $n^{th}$ cortical population |
| $C^n_{SST,BC}$ | 13.5 | SST to BC connectivity constant of $n^{th}$ cortical population |
| $C^n_{SST,VIP}$ | 20 | SST to VIP connectivity constant of $n^{th}$ cortical population |

| | | |
|---|---|---|
| $C^n_{VIP,SST}$ | 20 | VIP to SST connectivity constant of $n^{th}$ cortical population |
| $C^n_{BC}$ | 10 | BC to BC connectivity constant of $n^{th}$ cortical population |

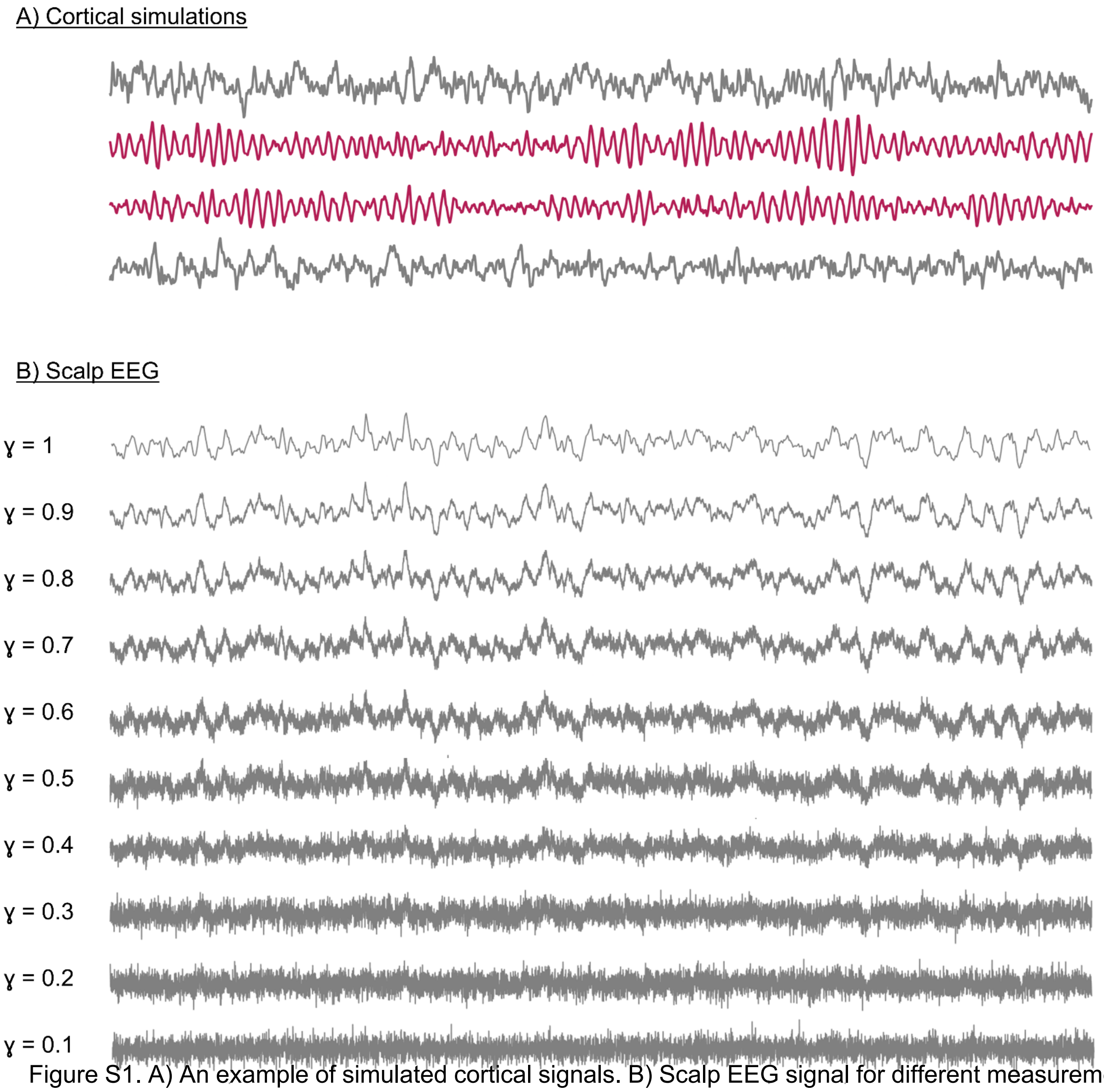


Figure S1. A) An example of simulated cortical signals. B) Scalp EEG signal for different measurement noise levels (i.e., different gamma values).

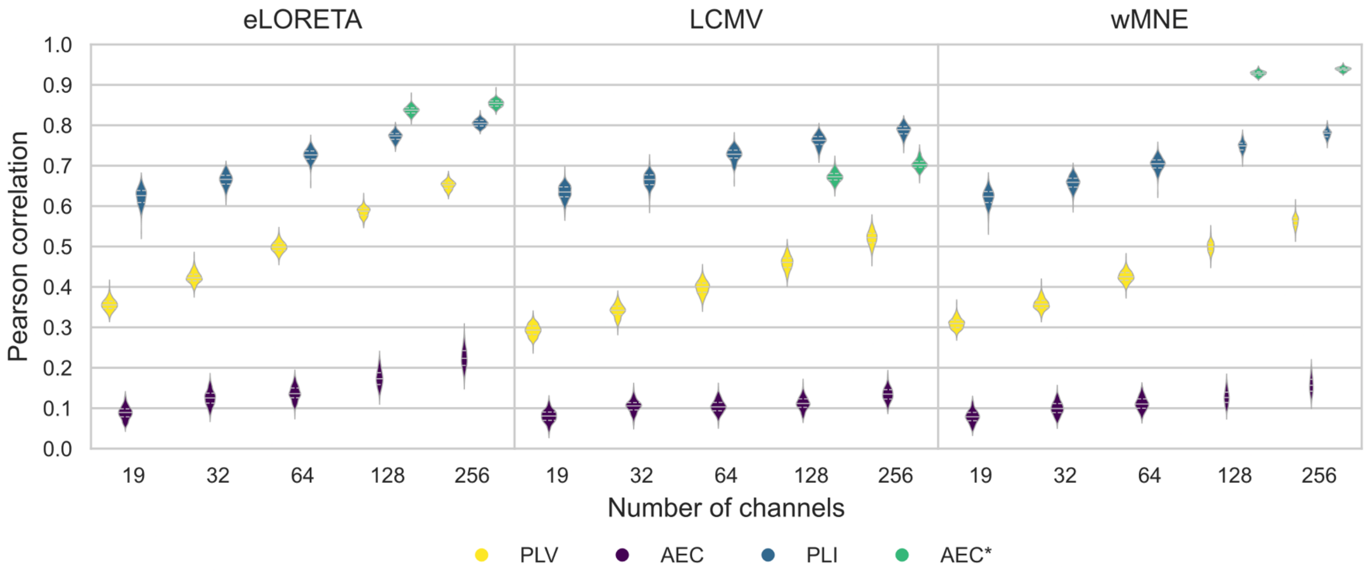


Figure S2. Violin plots of the Pearson correlation values computed between the reference and reconstructed DANs for all electrode montages and inverse methods/connectivity metrics combinations. *eLORETA - exact low resolution electromagnetic tomography. LCMV - linearly constrained minimum norm beamforming. wMNE - weighted minimum norm estimate. PLV - phase-locking value. AEC - amplitude envelope correlation. PLI - phase-lag index. AEC* - amplitude envelope correlation with source leakage correction.*

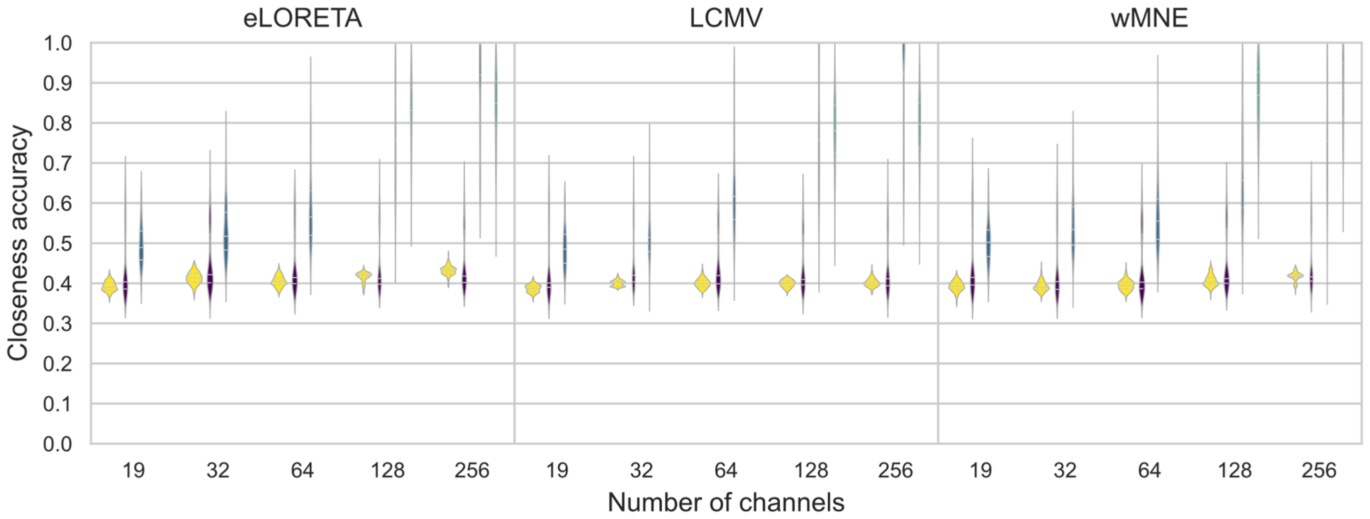


Figure S3. Violin plots of the Pearson correlation values computed between the reference and reconstructed DANs for all electrode montages and inverse methods/connectivity metrics combinations. *eLORETA - exact low resolution electromagnetic tomography. LCMV - linearly constrained minimum norm beamforming. wMNE - weighted minimum norm estimate. PLV - phase-locking value. AEC - amplitude envelope correlation. PLI - phase-lag index. AEC* - amplitude envelope correlation with source leakage correction.*

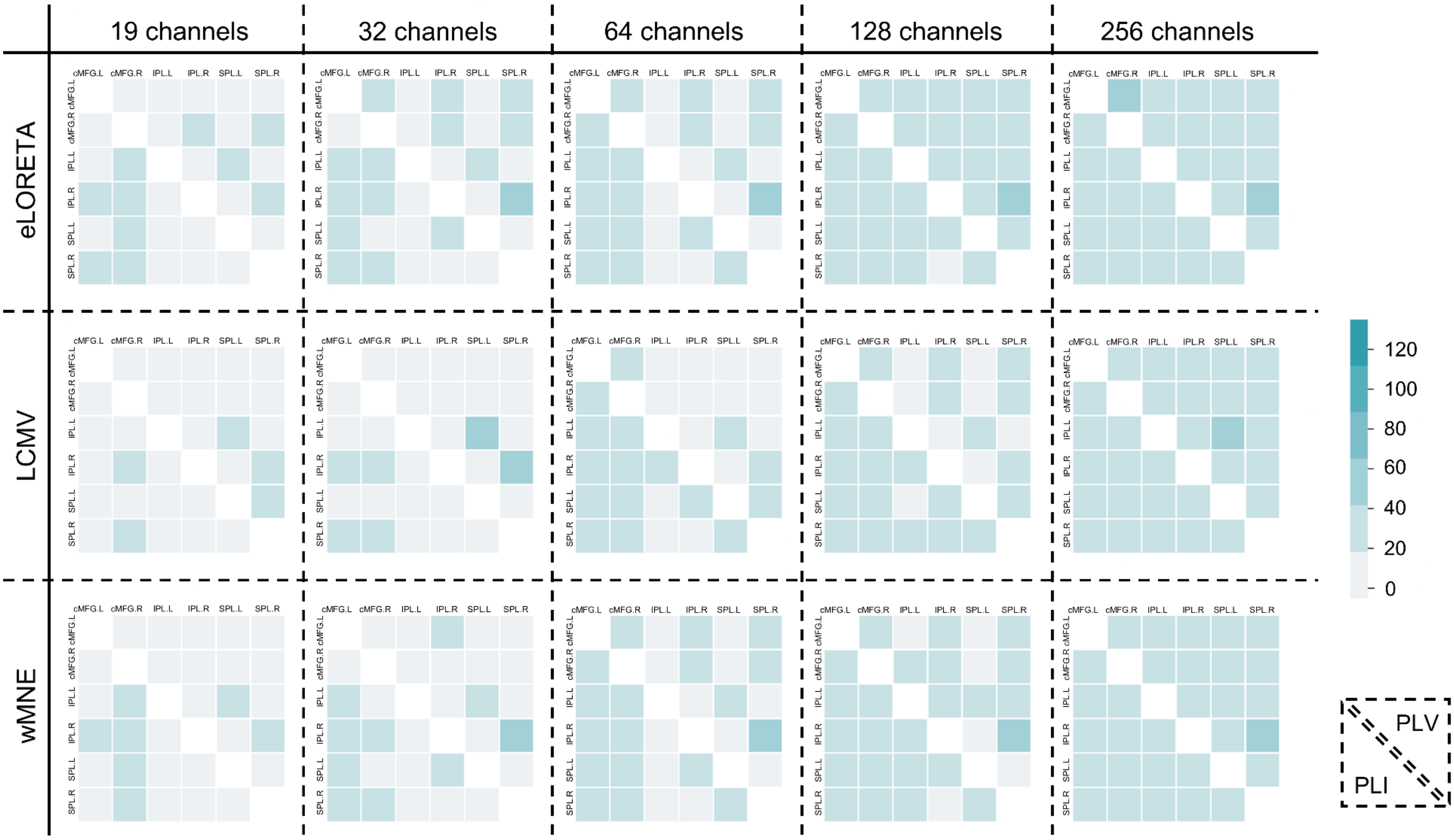

Figure S4. Heatmaps of contribution of the DAN edges (see Materials and Methods section) averaged across all subjects and epochs are shown for all sensor densities and inverse method/connectivity measure (PLV, PLI) combination. *eLORETA - exact low resolution electromagnetic tomography. LCMV - linearly constrained minimum norm beamforming. wMNE - weighted minimum norm estimate. PLV - phase-locking value. AEC - amplitude envelope correlation. PLI - phase-lag index. AEC* - amplitude envelope correlation with source leakage correction.*

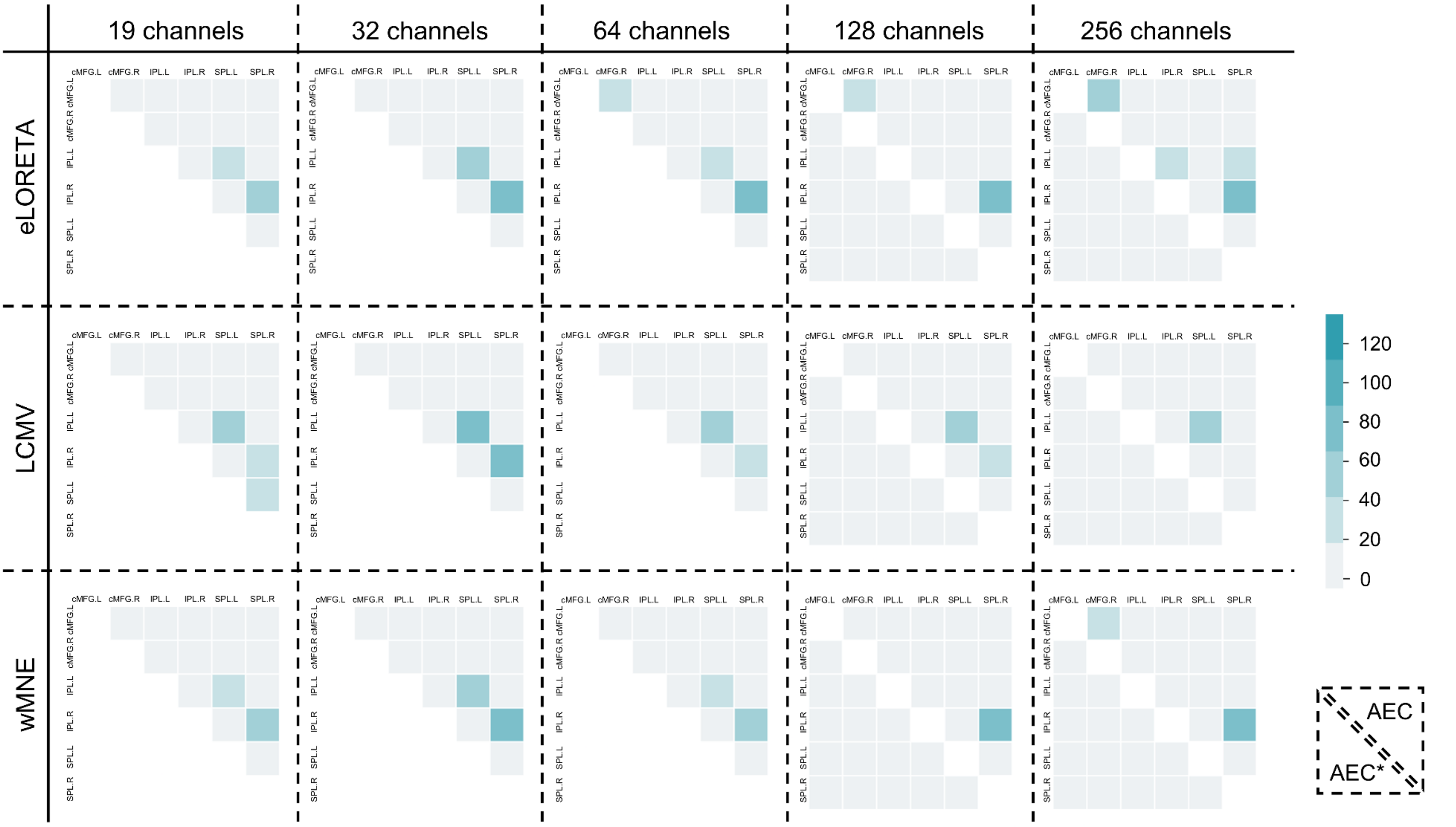


Figure S5. Heatmaps of contribution of the DAN edges averaged across all subjects and epochs are shown for all sensor densities and inverse method/connectivity measure (AEC, AEC*) combination. *eLORETA - exact low resolution electromagnetic tomography. LCMV - linearly constrained minimum norm beamforming. wMNE - weighted minimum norm estimate. PLV - phase-locking value. AEC - amplitude envelope correlation. PLI - phase-lag index. AEC* - amplitude envelope correlation with source leakage correction.*

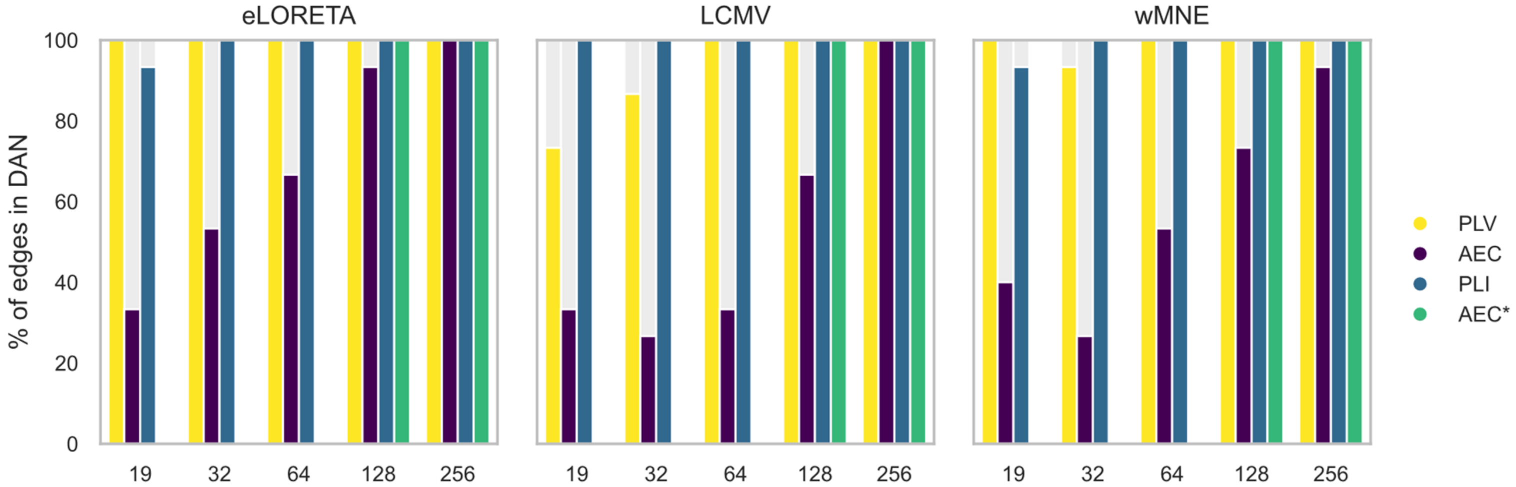


Figure S6. Barplots of the percentage of edges located within the DAN among the edges having the ≈ *0.7%* highest contribution values. *eLORETA - exact low resolution electromagnetic tomography. LCMV - linearly constrained minimum norm beamforming. wMNE - weighted minimum norm estimate. PLV - phase-locking value. AEC - amplitude envelope correlation. PLI - phase-lag index. AEC* - amplitude envelope correlation with source leakage correction.*

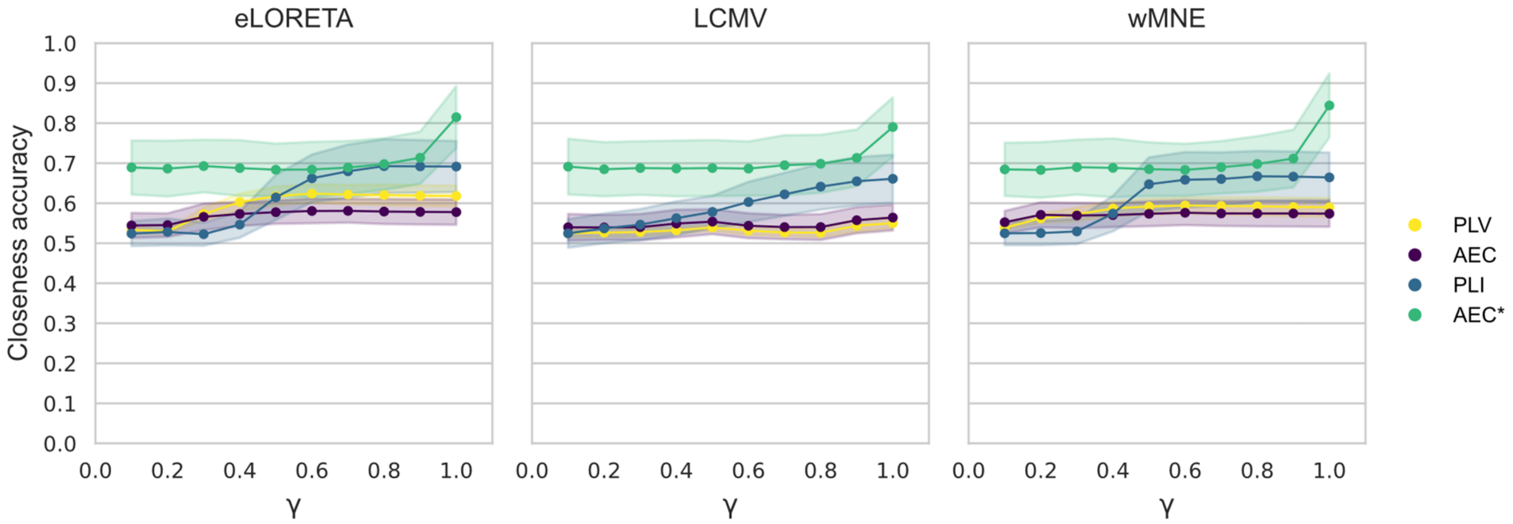


Figure S7. Mean and standard deviation of the closeness accuracy computed between the reference and reconstructed DMNs for different levels of measurement noise using 256 channels. *eLORETA - exact low resolution electromagnetic tomography. LCMV - linearly constrained minimum norm beamforming. wMNE - weighted minimum norm estimate. PLV - phase-locking value. AEC - amplitude envelope correlation. PLI - phase-lag index. AEC* - amplitude envelope correlation with source leakage correction.*

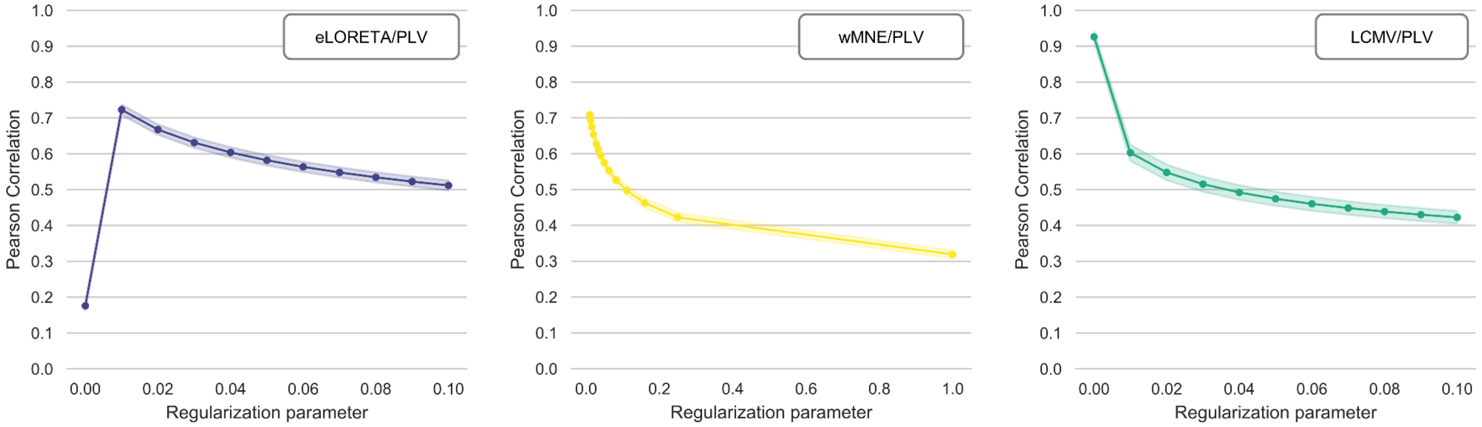


Figure S8. Mean and standard deviation of the Pearson correlation computed between the reference and reconstructed DMNs for different regularization values using 256 channels. *eLORETA - exact low resolution electromagnetic tomography. wMNE - weighted minimum norm estimate. LCMV - linearly constrained minimum norm beamforming. PLV - phase-locking value.*

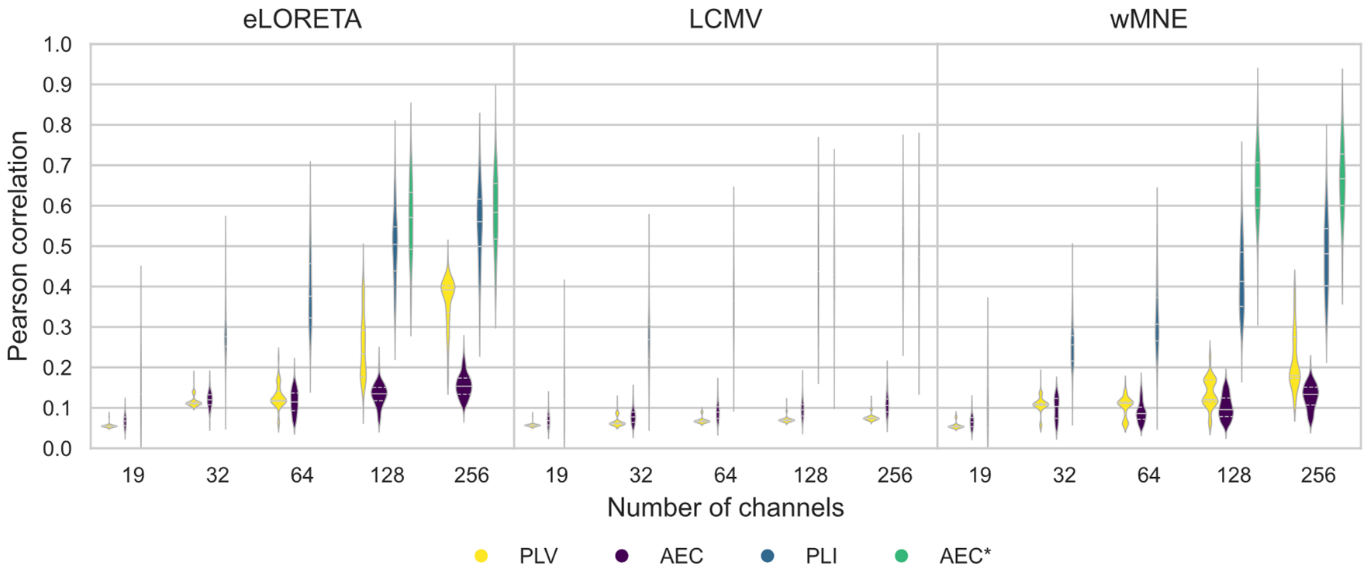


Figure S9. Violin plots of the Pearson correlation values computed between the reference and reconstructed DMNs (threshold = *1%*) for all electrode montages and inverse methods/connectivity metrics combinations. *eLORETA - exact low resolution electromagnetic tomography. LCMV - linearly constrained minimum norm beamforming. wMNE - weighted minimum norm estimate. PLV - phase-locking value. AEC - amplitude envelope correlation. PLI - phase-lag index. AEC* - amplitude envelope correlation with source leakage correction.*

### Statistical results

Tested parameters:

- Channels configurations (19, 32, 64, 128, 256)
- Inverse solutions (eLORETA, LCMV, wMNE)
- Connectivity measures (PLV, AEC, PLI)

**DMN - Pearson correlation**

| Factors | F | Pr(>Chisq) |
|---|---|---|
| Inverse method (2, 8906) | 4387.67 | 2.2e-16 *** |
| Connectivity measure (2, 8906) | 651671.77 | 2.2e-16 *** |
| Number of channels (4, 8906) | 32256.05 | 2.2e-16 *** |
| Inverse method × connectivity measure (4, 8096) | 547.74 | 2.2e-16 *** |
| Inverse method × channels (8, 8906) | 394 | 2.2e-16 *** |
| Connectivity measure × channels (8, 8906) | 1907 | 2.2e-16 *** |
| Inverse method × connectivity measure × channels (16, 8906) | 38.65 | 2.2e-16 *** |

R2m = 0.99
R2c = 0.99

| | Estimate | Std. Error | z-value | Pr(>\|z\|) |
|---|---|---|---|---|
| 32 - 19 == 0 | 0.035444 | 0.001687 | 21.01 | <2e-16 *** |
| 64 - 19 == 0 | 0.058163 | 0.001687 | 34.48 | <2e-16 *** |
| 128 - 19 == 0 | 0.102001 | 0.001687 | 60.47 | <2e-16 *** |
| 256 - 19 == 0 | 0.148868 | 0.001687 | 88.26 | <2e-16 *** |
| 64 - 32 == 0 | 0.022719 | 0.001687 | 13.47 | <2e-16 *** |
| 128 - 32 == 0 | 0.066557 | 0.001687 | 39.46 | <2e-16 *** |
| 256 - 32 == 0 | 0.113424 | 0.001687 | 67.25 | <2e-16 *** |
| 128 - 64 == 0 | 0.043839 | 0.001687 | 25.99 | <2e-16 *** |
| 256 - 64 == 0 | 0.090705 | 0.001687 | 53.78 | <2e-16 *** |
| 256 - 128 == 0 | 0.046867 | 0.001687 | 27.79 | <2e-16 *** |

| | Estimate | Std. Error | z-value | Pr(>\|z\|) |
|---|---|---|---|---|
| pli - aec == 0 | 0.456728 | 0.001687 | 270.8 | <2e-16 *** |
| plv - aec == 0 | 0.194675 | 0.001687 | 115.4 | <2e-16 *** |
| plv - pli == 0 | -0.262054 | 0.001687 | -155.4 | <2e-16 *** |

| | Estimate | Std. Error | z-value | Pr(>\|z\|) |
|---|---|---|---|---|
| lcmv - eloreta == 0 | 0.0072895 | 0.0016867 | 4.322 | <1e-04 *** |
| wmne - eloreta == 0 | 0.0001667 | 0.0016867 | 0.099 | 0.995 |
| wmne - lcmv == 0 | -0.0071229 | 0.0016867 | -4.223 | <1e-04 *** |

| | Estimate | Std. Error | z-value | Pr(>\|z\|) |
|---|---|---|---|---|
| lcmv.aec - eloreta.aec == 0 | 0.0072895 | 0.0016867 | 4.322 | <0.001 *** |
| wmne.aec - eloreta.aec == 0 | 0.0001667 | 0.0016867 | 0.099 | 1.000 |
| eloreta.pli - eloreta.aec == 0 | 0.4567283 | 0.0016867 | 270.777 | <0.001 *** |
| lcmv.pli - eloreta.aec == 0 | 0.4552020 | 0.0016867 | 269.873 | <0.001 *** |
| wmne.pli - eloreta.aec == 0 | 0.4463610 | 0.0016867 | 264.631 | <0.001 *** |
| eloreta.plv - eloreta.aec == 0 | 0.1946746 | 0.0016867 | 115.415 | <0.001 *** |
| lcmv.plv - eloreta.aec == 0 | 0.1911384 | 0.0016867 | 113.319 | <0.001 *** |
| wmne.plv - eloreta.aec == 0 | 0.1835499 | 0.0016867 | 108.820 | <0.001 *** |

| | | | | |
|---|---|---|---|---|
| wmne.aec - lcmv.aec == 0 | -0.0071229 | 0.0016867 | -4.223 | <0.001 *** |
| eloreta.pli - lcmv.aec == 0 | 0.4494387 | 0.0016867 | 266.456 | <0.001 *** |
| lcmv.pli - lcmv.aec == 0 | 0.4479125 | 0.0016867 | 265.551 | <0.001 *** |
| wmne.pli - lcmv.aec == 0 | 0.4390714 | 0.0016867 | 260.309 | <0.001 *** |
| eloreta.plv - lcmv.aec == 0 | 0.1873851 | 0.0016867 | 111.094 | <0.001 *** |
| lcmv.plv - lcmv.aec == 0 | 0.1838488 | 0.0016867 | 108.997 | <0.001 *** |
| wmne.plv - lcmv.aec == 0 | 0.1762604 | 0.0016867 | 104.498 | <0.001 *** |
| eloreta.pli - wmne.aec == 0 | 0.4565616 | 0.0016867 | 270.679 | <0.001 *** |
| lcmv.pli - wmne.aec == 0 | 0.4550353 | 0.0016867 | 269.774 | <0.001 *** |
| wmne.pli - wmne.aec == 0 | 0.4461943 | 0.0016867 | 264.532 | <0.001 *** |
| eloreta.plv - wmne.aec == 0 | 0.1945080 | 0.0016867 | 115.317 | <0.001 *** |
| lcmv.plv - wmne.aec == 0 | 0.1909717 | 0.0016867 | 113.220 | <0.001 *** |
| wmne.plv - wmne.aec == 0 | 0.1833832 | 0.0016867 | 108.721 | <0.001 *** |
| lcmv.pli - eloreta.pli == 0 | -0.0015263 | 0.0016867 | -0.905 | 0.993 |
| wmne.pli - eloreta.pli == 0 | -0.0103673 | 0.0016867 | -6.146 | <0.001 *** |
| eloreta.plv - eloreta.pli == 0 | -0.2620536 | 0.0016867 | -155.362 | <0.001 *** |
| lcmv.plv - eloreta.pli == 0 | -0.2655899 | 0.0016867 | -157.459 | <0.001 *** |
| wmne.plv - eloreta.pli == 0 | -0.2731783 | 0.0016867 | -161.957 | <0.001 *** |
| wmne.pli - lcmv.pli == 0 | -0.0088410 | 0.0016867 | -5.242 | <0.001 *** |
| eloreta.plv - lcmv.pli == 0 | -0.2605274 | 0.0016867 | -154.457 | <0.001 *** |
| lcmv.plv - lcmv.pli == 0 | -0.2640636 | 0.0016867 | -156.554 | <0.001 *** |
| wmne.plv - lcmv.pli == 0 | -0.2716521 | 0.0016867 | -161.053 | <0.001 *** |
| eloreta.plv - wmne.pli == 0 | -0.2516863 | 0.0016867 | -149.216 | <0.001 *** |
| lcmv.plv - wmne.pli == 0 | -0.2552226 | 0.0016867 | -151.312 | <0.001 *** |
| wmne.plv - wmne.pli == 0 | -0.2628110 | 0.0016867 | -155.811 | <0.001 *** |
| lcmv.plv - eloreta.plv == 0 | -0.0035363 | 0.0016867 | -2.097 | 0.476 |
| wmne.plv - eloreta.plv == 0 | -0.0111247 | 0.0016867 | -6.595 | <0.001 *** |
| wmne.plv - lcmv.plv == 0 | -0.0075884 | 0.0016867 | -4.499 | <0.001 *** |

**DMN - Closeness accuracy**

| Factors | F | Pr(>Chisq) |
|---|---|---|
| Inverse method (2,8906) | 304.21 | 2.2e-16 *** |
| Connectivity measure (2, 8906) | 2167.16 | 2.2e-16 *** |
| Number of channels (4, 8906) | 1278.20 | 2.2e-16 *** |
| Inverse method × connectivity measure (4, 8906) | 53.38 | 2.2e-16 *** |
| Inverse method × channels (8, 8906) | 83.60 | 2.2e-16 *** |
| Connectivity measure × channels (8, 8906) | 264.10 | 2.2e-16 *** |
| Inverse method × connectivity measure × channels (16, 8906) | 16.60 | 2.2e-16 *** |

R2m = 0.43
R2c = 0.44

| | **Estimate** | **Std. Error** | **z-value** | **Pr(>\|z\|)** |
|---|---|---|---|---|
| 32 - 19 == 0 | -0.113370 | 0.009587 | -11.825 | <0.001 *** |
| 64 - 19 == 0 | -0.133383 | 0.009587 | -13.912 | <0.001 *** |
| 128 - 19 == 0 | -0.143437 | 0.009587 | -14.961 | <0.001 *** |
| 256 - 19 == 0 | -0.136942 | 0.009587 | -14.283 | <0.001 *** |
| 64 - 32 == 0 | -0.020012 | 0.009587 | -2.087 | 0.2255 |
| 128 - 32 == 0 | -0.030067 | 0.009587 | -3.136 | 0.0148 * |
| 256 - 32 == 0 | -0.023571 | 0.009587 | -2.459 | 0.1002 |
| 128 - 64 == 0 | -0.010054 | 0.009587 | -1.049 | 0.8325 |

| | | | | |
|---|---|---|---|---|
| 256 - 64 == 0 | -0.003559 | 0.009587 | -0.371 | 0.9960 |
| 256 - 128 == 0 | 0.006495 | 0.009587 | 0.677 | 0.9613 |

| | **Estimate** | **Std. Error** | **z value** | **Pr(>\|z\|)** |
|---|---|---|---|---|
| pli - aec == 0 | 1.574e-05 | 9.587e-03 | 0.002 | 1 |
| plv - aec == 0 | 4.765e-02 | 9.587e-03 | 4.970 | 2.04e-06 *** |
| plv - pli == 0 | 4.763e-02 | 9.587e-03 | 4.968 | 1.82e-06 *** |

| | **Estimate** | **Std. Error** | **z-value** | **Pr(>\|z\|)** |
|---|---|---|---|---|
| lcmv - eloreta == 0 | -0.055200 | 0.009587 | -5.758 | <1e-04 *** |
| wmne - eloreta == 0 | -0.017196 | 0.009587 | -1.794 | 0.171723 |
| wmne - lcmv == 0 | 0.038004 | 0.009587 | 3.964 | 0.000204 *** |

| | **Estimate** | **Std. Error** | **z-value** | **Pr(>\|z\|)** |
|---|---|---|---|---|
| lcmv.aec - eloreta.aec == 0 | -5.520e-02 | 9.587e-03 | -5.758 | < 0.001 *** |
| wmne.aec - eloreta.aec == 0 | -1.720e-02 | 9.587e-03 | -1.794 | 0.68670 |
| eloreta.pli - eloreta.aec == 0 | 1.574e-05 | 9.587e-03 | 0.002 | 1.00000 |
| lcmv.pli - eloreta.aec == 0 | 1.211e-02 | 9.587e-03 | 1.263 | 0.94189 |
| wmne.pli - eloreta.aec == 0 | 2.526e-02 | 9.587e-03 | 2.634 | 0.17277 |
| eloreta.plv - eloreta.aec == 0 | 4.765e-02 | 9.587e-03 | 4.970 | < 0.001 *** |
| lcmv.plv - eloreta.aec == 0 | -2.202e-02 | 9.587e-03 | -2.297 | 0.34432 |
| wmne.plv - eloreta.aec == 0 | 1.785e-02 | 9.587e-03 | 1.861 | 0.64052 |
| wmne.aec - lcmv.aec == 0 | 3.800e-02 | 9.587e-03 | 3.964 | 0.00239 ** |
| eloreta.pli - lcmv.aec == 0 | 5.522e-02 | 9.587e-03 | 5.759 | < 0.001 *** |
| lcmv.pli - lcmv.aec == 0 | 6.731e-02 | 9.587e-03 | 7.020 | < 0.001 *** |
| wmne.pli - lcmv.aec == 0 | 8.046e-02 | 9.587e-03 | 8.392 | < 0.001 *** |
| eloreta.plv - lcmv.aec == 0 | 1.028e-01 | 9.587e-03 | 10.727 | < 0.001 *** |
| lcmv.plv - lcmv.aec == 0 | 3.318e-02 | 9.587e-03 | 3.461 | 0.01608 * |
| wmne.plv - lcmv.aec == 0 | 7.304e-02 | 9.587e-03 | 7.619 | < 0.001 *** |
| eloreta.pli - wmne.aec == 0 | 1.721e-02 | 9.587e-03 | 1.795 | 0.68572 |
| lcmv.pli - wmne.aec == 0 | 2.930e-02 | 9.587e-03 | 3.057 | 0.05744 . |
| wmne.pli - wmne.aec == 0 | 4.245e-02 | 9.587e-03 | 4.428 | < 0.001 *** |
| eloreta.plv - wmne.aec == 0 | 6.484e-02 | 9.587e-03 | 6.763 | < 0.001 *** |
| lcmv.plv - wmne.aec == 0 | -4.823e-03 | 9.587e-03 | -0.503 | 0.99990 |
| wmne.plv - wmne.aec == 0 | 3.504e-02 | 9.587e-03 | 3.655 | 0.00772 ** |
| lcmv.pli - eloreta.pli == 0 | 1.209e-02 | 9.587e-03 | 1.261 | 0.94233 |
| wmne.pli - eloreta.pli == 0 | 2.524e-02 | 9.587e-03 | 2.633 | 0.17285 |
| eloreta.plv - eloreta.pli == 0 | 4.763e-02 | 9.587e-03 | 4.968 | < 0.001 *** |
| lcmv.plv - eloreta.pli == 0 | -2.203e-02 | 9.587e-03 | -2.298 | 0.34285 |
| wmne.plv - eloreta.pli == 0 | 1.783e-02 | 9.587e-03 | 1.860 | 0.64187 |
| wmne.pli - lcmv.pli == 0 | 1.315e-02 | 9.587e-03 | 1.372 | 0.90866 |
| eloreta.plv - lcmv.pli == 0 | 3.554e-02 | 9.587e-03 | 3.707 | 0.00635 ** |
| lcmv.plv - lcmv.pli == 0 | -3.413e-02 | 9.587e-03 | -3.560 | 0.01132 * |
| wmne.plv - lcmv.pli == 0 | 5.737e-03 | 9.587e-03 | 0.598 | 0.99962 |
| eloreta.plv - wmne.pli == 0 | 2.239e-02 | 9.587e-03 | 2.335 | 0.32077 |
| lcmv.plv - wmne.pli == 0 | -4.728e-02 | 9.587e-03 | -4.931 | < 0.001 *** |
| wmne.plv - wmne.pli == 0 | -7.412e-03 | 9.587e-03 | -0.773 | 0.99756 |
| lcmv.plv - eloreta.plv == 0 | -6.967e-02 | 9.587e-03 | -7.266 | < 0.001 *** |
| wmne.plv - eloreta.plv == 0 | -2.980e-02 | 9.587e-03 | -3.108 | 0.04896 * |
| wmne.plv - lcmv.plv == 0 | 3.986e-02 | 9.587e-03 | 4.158 | 0.00101 ** |

**DAN - Pearson correlation**

| Factors | F | Pr(>Chisq) |
|---|---|---|
| Inverse method (2,8906) | 8649.7 | 2.2e-16 *** |
| Connectivity measure (2,8906) | 1088000 | 2.2e-16 *** |
| Number of channels (4, 8906) | 34942 | 2.2e-16 *** |

| | | |
|---|---|---|
| Inverse method × connectivity measure (8, 8906) | 2538.8 | 2.2e-16 *** |
| Inverse method × channels (8, 8906) | 373.98 | 2.2e-16 *** |
| Connectivity measure × channels (8, 8906) | 3344.9 | 2.2e-16 *** |
| Inverse method × connectivity measure × channels (16, 8906) | 39.522 | 2.2e-16 *** |

R2m = 0.9959
R2c = 0.9962

| | Estimate | Std. Error | z-value | Pr(>\|z\|) |
|---|---|---|---|---|
| 32 - 19 == 0 | 0.036506 | 0.001546 | 23.62 | <1e-10 *** |
| 64 - 19 == 0 | 0.048300 | 0.001546 | 31.25 | <1e-10 *** |
| 128 - 19 == 0 | 0.085999 | 0.001546 | 55.64 | <1e-10 *** |
| 256 - 19 == 0 | 0.135554 | 0.001546 | 87.70 | <1e-10 *** |
| 64 - 32 == 0 | 0.011794 | 0.001546 | 7.63 | <1e-10 *** |
| 128 - 32 == 0 | 0.049493 | 0.001546 | 32.02 | <1e-10 *** |
| 256 - 32 == 0 | 0.099048 | 0.001546 | 64.08 | <1e-10 *** |
| 128 - 64 == 0 | 0.037699 | 0.001546 | 24.39 | <1e-10 *** |
| 256 - 64 == 0 | 0.087255 | 0.001546 | 56.45 | <1e-10 *** |
| 256 - 128 == 0 | 0.049555 | 0.001546 | 32.06 | <1e-10 *** |

| | Estimate | Std. Error | z-value | Pr(>\|z\|) |
|---|---|---|---|---|
| pli - aec == 0 | 0.534939 | 0.001546 | 346.1 | <2e-16 *** |
| plv - aec == 0 | 0.267440 | 0.001546 | 173.0 | <2e-16 *** |
| plv - pli == 0 | -0.267498 | 0.001546 | -173.1 | <2e-16 *** |

| | Estimate | Std. Error | z-value | Pr(>\|z\|) |
|---|---|---|---|---|
| lcmv - eloreta == 0 | -0.0082866 | 0.0015456 | -5.361 | <1e-04 *** |
| wmne - eloreta == 0 | -0.0092315 | 0.0015456 | -5.973 | <1e-04 *** |
| wmne - lcmv == 0 | -0.0009449 | 0.0015456 | -0.611 | 0.814 |

| | Estimate | Std. Error | z-value | Pr(>\|z\|) |
|---|---|---|---|---|
| lcmv.aec - eloreta.aec == 0 | -0.0082866 | 0.0015456 | -5.361 | <1e-04 *** |
| wmne.aec - eloreta.aec == 0 | -0.0092315 | 0.0015456 | -5.973 | <1e-04 *** |
| eloreta.pli - eloreta.aec == 0 | 0.5349388 | 0.0015456 | 346.094 | <1e-04 *** |
| lcmv.pli - eloreta.aec == 0 | 0.5454049 | 0.0015456 | 352.865 | <1e-04 *** |
| wmne.pli - eloreta.aec == 0 | 0.5331581 | 0.0015456 | 344.942 | <1e-04 *** |
| eloreta.plv - eloreta.aec == 0 | 0.2674405 | 0.0015456 | 173.028 | <1e-04 *** |
| lcmv.plv - eloreta.aec == 0 | 0.2038661 | 0.0015456 | 131.897 | <1e-04 *** |
| wmne.plv - eloreta.aec == 0 | 0.2210464 | 0.0015456 | 143.012 | <1e-04 *** |
| wmne.aec - lcmv.aec == 0 | -0.0009449 | 0.0015456 | -0.611 | 1.000 |
| eloreta.pli - lcmv.aec == 0 | 0.5432254 | 0.0015456 | 351.455 | <1e-04 *** |
| lcmv.pli - lcmv.aec == 0 | 0.5536915 | 0.0015456 | 358.227 | <1e-04 *** |
| wmne.pli - lcmv.aec == 0 | 0.5414447 | 0.0015456 | 350.303 | <1e-04 *** |
| eloreta.plv - lcmv.aec == 0 | 0.2757271 | 0.0015456 | 178.390 | <1e-04 *** |
| lcmv.plv - lcmv.aec == 0 | 0.2121527 | 0.0015456 | 137.258 | <1e-04 *** |
| wmne.plv - lcmv.aec == 0 | 0.2293330 | 0.0015456 | 148.374 | <1e-04 *** |
| eloreta.pli - wmne.aec == 0 | 0.5441703 | 0.0015456 | 352.067 | <1e-04 *** |
| lcmv.pli - wmne.aec == 0 | 0.5546364 | 0.0015456 | 358.838 | <1e-04 *** |
| wmne.pli - wmne.aec == 0 | 0.5423896 | 0.0015456 | 350.914 | <1e-04 *** |
| eloreta.plv - wmne.aec == 0 | 0.2766720 | 0.0015456 | 179.001 | <1e-04 *** |
| lcmv.plv - wmne.aec == 0 | 0.2130976 | 0.0015456 | 137.870 | <1e-04 *** |
| wmne.plv - wmne.aec == 0 | 0.2302778 | 0.0015456 | 148.985 | <1e-04 *** |
| lcmv.pli - eloreta.pli == 0 | 0.0104661 | 0.0015456 | 6.771 | <1e-04 *** |

| | | | | |
|---|---|---|---|---|
| wmne.pli - eloreta.pli == 0 | -0.0017807 | 0.0015456 | -1.152 | 0.966 |
| eloreta.plv - eloreta.pli == 0 | -0.2674983 | 0.0015456 | -173.066 | <1e-04 *** |
| lcmv.plv - eloreta.pli == 0 | -0.3310727 | 0.0015456 | -214.197 | <1e-04 *** |
| wmne.plv - eloreta.pli == 0 | -0.3138924 | 0.0015456 | -203.082 | <1e-04 *** |
| wmne.pli - lcmv.pli == 0 | -0.0122468 | 0.0015456 | -7.923 | <1e-04 *** |
| eloreta.plv - lcmv.pli == 0 | -0.2779644 | 0.0015456 | -179.837 | <1e-04 *** |
| lcmv.plv - lcmv.pli == 0 | -0.3415388 | 0.0015456 | -220.968 | <1e-04 *** |
| wmne.plv - lcmv.pli == 0 | -0.3243585 | 0.0015456 | -209.853 | <1e-04 *** |
| eloreta.plv - wmne.pli == 0 | -0.2657176 | 0.0015456 | -171.914 | <1e-04 *** |
| lcmv.plv - wmne.pli == 0 | -0.3292920 | 0.0015456 | -213.045 | <1e-04 *** |
| wmne.plv - wmne.pli == 0 | -0.3121117 | 0.0015456 | -201.930 | <1e-04 *** |
| lcmv.plv - eloreta.plv == 0 | -0.0635744 | 0.0015456 | -41.131 | <1e-04 *** |
| wmne.plv - eloreta.plv == 0 | -0.0463941 | 0.0015456 | -30.016 | <1e-04 *** |
| wmne.plv - lcmv.plv == 0 | 0.0171803 | 0.0015456 | 11.115 | <1e-04 *** |

**DAN - Closeness accuracy**

| Factors | F | Pr(>Chisq) |
|---|---|---|
| Inverse method (2,8906) | 25.4760 | 9.278e-12 *** |
| Connectivity measure (2,8906) | 9729.6266 | < 2.2e-16 *** |
| Number of channels (4, 8906) | 564.1601 | < 2.2e-16 *** |
| Inverse method × connectivity measure (2, 8906) | 18.6607 | 2.748e-15 *** |
| Inverse method × channels (8, 8906) | 10.2845 | 2.001e-14 *** |
| Connectivity measure × channels (8, 8906) | 326.9663 | 2.2e-16 *** |
| Inverse method × connectivity measure × channels (16, 8906) | 9.6136 | 2.2e-16 *** |

R2m = 0.72
R2c = 0.74

| | **Estimate** | **Std. Error** | **z-value** | **Pr(>\|z\|)** |
|---|---|---|---|---|
| 32 - 19 == 0 | -0.0604201 | 0.0237203 | -2.547 | 0.0805 . |
| 64 - 19 == 0 | -0.0521371 | 0.0237203 | -2.198 | 0.1803 |
| 128 - 19 == 0 | -0.0517862 | 0.0237203 | -2.183 | 0.1859 |
| 256 - 19 == 0 | -0.0645695 | 0.0237203 | -2.722 | 0.0507 . |
| 64 - 32 == 0 | 0.0082830 | 0.0237203 | 0.349 | 0.9968 |
| 128 - 32 == 0 | 0.0086338 | 0.0237203 | 0.364 | 0.9963 |
| 256 - 32 == 0 | -0.0041495 | 0.0237203 | -0.175 | 0.9998 |
| 128 - 64 == 0 | 0.0003509 | 0.0237203 | 0.015 | 1.0000 |
| 256 - 64 == 0 | -0.0124324 | 0.0237203 | -0.524 | 0.9849 |
| 256 - 128 == 0 | -0.0127833 | 0.0237203 | -0.539 | 0.9833 |

| | **Estimate** | **Std. Error** | **z-value** | **Pr(>\|z\|)** |
|---|---|---|---|---|
| pli - aec == 0 | -0.27054 | 0.02372 | -11.40 | <2e-16 *** |
| plv - aec == 0 | 0.25018 | 0.02372 | 10.55 | <2e-16 *** |
| plv - pli == 0 | 0.52072 | 0.02372 | 21.95 | <2e-16 *** |

| | **Estimate** | **Std. Error** | **z-value** | **Pr(>\|z\|)** |
|---|---|---|---|---|
| lcmv - eloreta == 0 | -0.001967 | 0.023720 | -0.083 | 0.9962 |
| wmne - eloreta == 0 | -0.057405 | 0.023720 | -2.420 | 0.0411 * |
| wmne - lcmv == 0 | -0.055438 | 0.023720 | -2.337 | 0.0508 . |

| | Estimate | Std. Error | z-value | Pr(>\|z\|) |
|---|---|---|---|---|
| lcmv.aec - eloreta.aec == 0 | -0.001967 | 0.023720 | -0.083 | 1.000 |
| wmne.aec - eloreta.aec == 0 | -0.057405 | 0.023720 | -2.420 | 0.273 |
| eloreta.pli - eloreta.aec == 0 | -0.270535 | 0.023720 | -11.405 | <0.001 *** |
| lcmv.pli - eloreta.aec == 0 | -0.236337 | 0.023720 | -9.964 | <0.001 *** |
| wmne.pli - eloreta.aec == 0 | -0.302323 | 0.023720 | -12.745 | <0.001 *** |
| eloreta.plv - eloreta.aec == 0 | 0.250185 | 0.023720 | 10.547 | <0.001 *** |
| lcmv.plv - eloreta.aec == 0 | 0.283550 | 0.023720 | 11.954 | <0.001 *** |
| wmne.plv - eloreta.aec == 0 | 0.248732 | 0.023720 | 10.486 | <0.001 *** |
| wmne.aec - lcmv.aec == 0 | -0.055438 | 0.023720 | -2.337 | 0.320 |
| eloreta.pli - lcmv.aec == 0 | 0.268569 | 0.023720 | -11.322 | <0.001 *** |
| lcmv.pli - lcmv.aec == 0 | -0.234371 | 0.023720 | -9.881 | <0.001 *** |
| wmne.pli - lcmv.aec == 0 | -0.300356 | 0.023720 | -12.662 | <0.001 *** |
| eloreta.plv - lcmv.aec == 0 | 0.252151 | 0.023720 | 10.630 | <0.001 *** |
| lcmv.plv - lcmv.aec == 0 | 0.285516 | 0.023720 | 12.037 | <0.001 *** |
| wmne.plv - lcmv.aec == 0 | 0.250699 | 0.023720 | 10.569 | <0.001 *** |
| eloreta.pli - wmne.aec == 0 | -0.213131 | 0.023720 | -8.985 | <0.001 *** |
| lcmv.pli - wmne.aec == 0 | -0.178933 | 0.023720 | -7.543 | <0.001 *** |
| wmne.pli - wmne.aec == 0 | -0.244918 | 0.023720 | -10.325 | <0.001 *** |
| eloreta.plv - wmne.aec == 0 | 0.307589 | 0.023720 | 12.967 | <0.001 *** |
| lcmv.plv - wmne.aec == 0 | 0.340954 | 0.023720 | 14.374 | <0.001 *** |
| wmne.plv - wmne.aec == 0 | 0.306137 | 0.023720 | 12.906 | <0.001 *** |
| lcmv.pli - eloreta.pli == 0 | 0.034198 | 0.023720 | 1.442 | 0.882 |
| wmne.pli - eloreta.pli == 0 | -0.031788 | 0.023720 | -1.340 | 0.919 |
| eloreta.plv - eloreta.pli == 0 | 0.520720 | .023720 | 21.953 | <0.001 *** |
| lcmv.plv - eloreta.pli == 0 | 0.554085 | 0.023720 | 23.359 | <0.001 *** |
| wmne.plv - eloreta.pli == 0 | 0.519267 | 0.023720 | 21.891 | <0.001 *** |
| wmne.pli - lcmv.pli == 0 | -0.065986 | 0.023720 | -2.782 | 0.121 |
| eloreta.plv - lcmv.pli == 0 | 0.486522 | 0.023720 | 20.511 | <0.001 *** |
| lcmv.plv - lcmv.pli == 0 | 0.519887 | 0.023720 | 21.917 | <0.001 *** |
| wmne.plv - lcmv.pli == 0 | 0.485069 | 0.023720 | 20.450 | <0.001 |
| eloreta.plv - wmne.pli == 0 | 0.552508 | 0.023720 | 23.293 | <0.001 *** |
| lcmv.plv - wmne.pli == 0 | 0.585873 | 0.023720 | 24.699 | <0.001 *** |
| wmne.plv - wmne.pli == 0 | 0.551055 | 0.023720 | 23.231 | <0.001 *** |
| lcmv.plv - eloreta.plv == 0 | 0.033365 | 0.023720 | 1.407 | 0.896 |
| wmne.plv - eloreta.plv == 0 | -0.001453 | 0.023720 | -0.061 | 1.000 |
| wmne.plv - lcmv.plv == 0 | -0.034818 | 0.023720 | -1.468 | 0.871 |

### Statistical results

Tested parameters:

- Channels configurations (128, 256)
- Inverse solutions (eLORETA, LCMV, wMNE)
- Connectivity measures (PLV, AEC, PLI, AEC*)

(N.B.: AEC* == aec_orth)

**DMN - Pearson correlation**

| Factors | F | Pr(>Chisq) |
|---|---|---|
| Inverse method (2, 4727) | 10396.12 | 2.2e-16 *** |
| Connectivity measure (3, 4727) | 371597.94 | 2.2e-16 *** |
| Number of channels (1, 8906) | 6142.34 | 2.2e-16 *** |
| Inverse method × connectivity measure (6, 4727) | 4858.35 | 2.2e-16 *** |
| Inverse method × channels (2, 4727) | 19.98 | 2.2e-16 *** |
| Connectivity measure × channels (3, 4727) | 424.18 | 2.2e-16 *** |
| Inverse method × connectivity measure × channels (6, 4727) | 24.13 | 2.2e-16 *** |

R2m = 0.9956
R2c = 0.9959

| | **Estimate** | **Std. Error** | **z-value** | **Pr(>\|z\|)** |
|---|---|---|---|---|
| aec_orth - aec == 0 | 0.655102 | 0.001604 | 408.52 | <2e-16 *** |
| pli - aec == 0 | 0.519730 | 0.001604 | 324.10 | <2e-16 *** |
| plv - aec == 0 | 0.341390 | 0.001604 | 212.89 | <2e-16 *** |
| pli - aec_orth == 0 | -0.135372 | 0.001604 | -84.42 | <2e-16 *** |
| plv - aec_orth == 0 | -0.313712 | 0.001604 | -195.63 | <2e-16 *** |
| plv - pli == 0 | -0.178340 | 0.001604 | -111.21 | <2e-16 *** |

| | **Estimate** | **Std. Error** | **z-value** | **Pr(>\|z\|)** |
|---|---|---|---|---|
| lcmv - eloreta == 0 | -0.034123 | 0.001604 | -21.28 | <0.001 *** |
| wmne - eloreta == 0 | -0.038822 | 0.001604 | -24.21 | <0.001 *** |
| wmne - lcmv == 0 | -0.004699 | 0.001604 | -2.93 | 0.0095 ** |

| | **Estimate** | **Std. Error** | **z-value** | **Pr(>\|z\|)** |
|---|---|---|---|---|
| lcmv.aec - eloreta.aec == 0 | -0.034123 | 0.001604 | -21.279 | <0.001 *** |
| wmne.aec - eloreta.aec == 0 | -0.038822 | 0.001604 | -24.209 | <0.001 *** |
| eloreta.aec_orth - eloreta.aec == 0 | 0.655102 | 0.001604 | 408.517 | <0.001 *** |
| lcmv.aec_orth - eloreta.aec == 0 | 0.510187 | 0.001604 | 318.149 | <0.001 *** |
| wmne.aec_orth - eloreta.aec == 0 | 0.738714 | 0.001604 | 460.657 | <0.001 *** |
| eloreta.pli - eloreta.aec == 0 | 0.519730 | 0.001604 | 324.100 | <0.001 *** |
| lcmv.pli - eloreta.aec == 0 | 0.487668 | 0.001604 | 304.106 | <0.001 *** |
| wmne.pli - eloreta.aec == 0 | 0.485484 | 0.001604 | 302.744 | <0.001 *** |
| eloreta.plv - eloreta.aec == 0 | 0.341390 | 0.001604 | 212.888 | <0.001 *** |
| lcmv.plv - eloreta.aec == 0 | 0.246996 | 0.001604 | 154.025 | <0.001 *** |
| wmne.plv - eloreta.aec == 0 | 0.263516 | 0.001604 | 164.327 | <0.001 *** |
| wmne.aec - lcmv.aec == 0 | -0.004699 | 0.001604 | -2.930 | 0.131 |
| eloreta.aec_orth - lcmv.aec == 0 | 0.689226 | 0.001604 | 429.796 | <0.001 *** |
| lcmv.aec_orth - lcmv.aec == 0 | 0.544311 | 0.001604 | 339.428 | <0.001 *** |
| wmne.aec_orth - lcmv.aec == 0 | 0.772838 | 0.001604 | 481.936 | <0.001 *** |

| | | | | |
|---|---|---|---|---|
| eloreta.pli - lcmv.aec == 0 | 0.553853 | 0.001604 | 345.379 | <0.001 *** |
| lcmv.pli - lcmv.aec == 0 | 0.521791 | 0.001604 | 325.385 | <0.001 *** |
| wmne.pli - lcmv.aec == 0 | 0.519608 | 0.001604 | 324.023 | <0.001 *** |
| eloreta.plv - lcmv.aec == 0 | 0.375513 | 0.001604 | 234.167 | <0.001 *** |
| lcmv.plv - lcmv.aec == 0 | 0.281119 | 0.001604 | 175.304 | <0.001 *** |
| wmne.plv - lcmv.aec == 0 | 0.297639 | 0.001604 | 185.606 | <0.001 *** |
| eloreta.aec_orth - wmne.aec == 0 | 0.693924 | 0.001604 | 432.726 | <0.001 *** |
| lcmv.aec_orth - wmne.aec == 0 | 0.549009 | 0.001604 | 342.358 | <0.001 *** |
| wmne.aec_orth - wmne.aec == 0 | 0.777536 | 0.001604 | 484.866 | <0.001 *** |
| eloreta.pli - wmne.aec == 0 | 0.558552 | 0.001604 | 348.309 | <0.001 *** |
| lcmv.pli - wmne.aec == 0 | 0.526490 | 0.001604 | 328.315 | <0.001 *** |
| wmne.pli - wmne.aec == 0 | 0.524306 | 0.001604 | 326.954 | <0.001 *** |
| eloreta.plv - wmne.aec == 0 | 0.380212 | 0.001604 | 237.097 | <0.001 *** |
| lcmv.plv - wmne.aec == 0 | 0.285818 | 0.001604 | 178.234 | <0.001 *** |
| wmne.plv - wmne.aec == 0 | 0.302338 | 0.001604 | 188.536 | <0.001 *** |
| lcmv.aec_orth - eloreta.aec_orth == 0 | -0.144915 | 0.001604 | -90.368 | <0.001 *** |
| wmne.aec_orth - eloreta.aec_orth == 0 | 0.083612 | 0.001604 | 52.140 | <0.001 *** |
| eloreta.pli - eloreta.aec_orth == 0 | -0.135372 | 0.001604 | -84.417 | <0.001 *** |
| lcmv.pli - eloreta.aec_orth == 0 | -0.167434 | 0.001604 | -104.411 | <0.001 *** |
| wmne.pli - eloreta.aec_orth == 0 | -0.169618 | 0.001604 | -105.772 | <0.001 *** |
| eloreta.plv - eloreta.aec_orth == 0 | -0.313712 | 0.001604 | -195.629 | <0.001 *** |
| lcmv.plv - eloreta.aec_orth == 0 | -0.408106 | 0.001604 | -254.492 | <0.001 *** |
| wmne.plv - eloreta.aec_orth == 0 | -0.391586 | 0.001604 | -244.190 | <0.001 *** |
| wmne.aec_orth - lcmv.aec_orth == 0 | 0.228527 | 0.001604 | 142.508 | <0.001 *** |
| eloreta.pli - lcmv.aec_orth == 0 | 0.009543 | 0.001604 | 5.951 | <0.001 *** |
| lcmv.pli - lcmv.aec_orth == 0 | -0.022519 | 0.001604 | -14.043 | <0.001 *** |
| wmne.pli - lcmv.aec_orth == 0 | -0.024703 | 0.001604 | -15.405 | <0.001 *** |
| eloreta.plv - lcmv.aec_orth == 0 | -0.168798 | 0.001604 | -105.261 | <0.001 *** |
| lcmv.plv - lcmv.aec_orth == 0 | -0.263192 | 0.001604 | -164.124 | <0.001 *** |
| wmne.plv - lcmv.aec_orth == 0 | -0.246671 | 0.001604 | -153.822 | <0.001 *** |
| eloreta.pli - wmne.aec_orth == 0 | -0.218984 | 0.001604 | -136.557 | <0.001 *** |
| lcmv.pli - wmne.aec_orth == 0 | -0.251046 | 0.001604 | -156.551 | <0.001 *** |
| wmne.pli - wmne.aec_orth == 0 | -0.253230 | 0.001604 | -157.912 | <0.001 *** |
| eloreta.plv - wmne.aec_orth == 0 | -0.397325 | 0.001604 | -247.769 | <0.001 *** |
| lcmv.plv - wmne.aec_orth == 0 | -0.491719 | 0.001604 | -306.632 | <0.001 *** |
| wmne.plv - wmne.aec_orth == 0 | -0.475198 | 0.001604 | -296.330 | <0.001 *** |
| lcmv.pli - eloreta.pli == 0 | -0.032062 | 0.001604 | -19.994 | <0.001 *** |
| wmne.pli - eloreta.pli == 0 | -0.034246 | 0.001604 | -21.355 | <0.001 *** |
| eloreta.plv - eloreta.pli == 0 | -0.178340 | 0.001604 | -111.212 | <0.001 *** |
| lcmv.plv - eloreta.pli == 0 | -0.272734 | 0.001604 | -170.075 | <0.001 *** |
| wmne.plv - eloreta.pli == 0 | -0.256214 | 0.001604 | -159.773 | <0.001 *** |
| wmne.pli - lcmv.pli == 0 | -0.002184 | 0.001604 | -1.362 | 0.970 |
| eloreta.plv - lcmv.pli == 0 | -0.146278 | 0.001604 | -91.218 | <0.001 *** |
| lcmv.plv - lcmv.pli == 0 | -0.240672 | 0.001604 | -150.081 | <0.001 *** |
| wmne.plv - lcmv.pli == 0 | -0.224152 | 0.001604 | -139.779 | <0.001 *** |
| eloreta.plv - wmne.pli == 0 | -0.144094 | 0.001604 | -89.856 | <0.001 *** |
| lcmv.plv - wmne.pli == 0 | -0.238489 | 0.001604 | -148.720 | <0.001 *** |
| wmne.plv - wmne.pli == 0 | -0.221968 | 0.001604 | -138.418 | <0.001 *** |
| lcmv.plv - eloreta.plv == 0 | -0.094394 | 0.001604 | -58.863 | <0.001 *** |
| wmne.plv - eloreta.plv == 0 | -0.077874 | 0.001604 | -48.561 | <0.001 *** |
| wmne.plv - lcmv.plv == 0 | 0.016520 | 0.001604 | 10.302 | <0.001 *** |

##### DMN - Closeness accuracy

| Factors | F | Pr(>Chisq) |
|---|---|---|
| Inverse method (2, 4727) | 394.91 | 2.2e-16 *** |

| | | |
|---|---|---|
| Connectivity measure (3, 4727) | 6696.08 | 2.2e-16 *** |
| Number of channels (1, 4727) | 267.44 | 2.2e-16 *** |
| Inverse method × connectivity measure (6, 4727) | 74.41 | 2.2e-16 *** |
| Inverse method × channels (2, 4727) | 11.05 | 2.2e-16 *** |
| Connectivity measure × channels (3, 4727) | 20.91 | 2.2e-16 *** |
| Inverse method × connectivity measure × channels (6, 4727) | 5.31 | 2.2e-16 *** |

R2m = 0.81
R2c = 0.82

| | Estimate | Std. Error | z-value | Pr(>\|z\|) |
|---|---|---|---|---|
| aec_orth - aec == 0 | -0.47856 | 0.01050 | -45.580 | <1e-05 *** |
| pli - aec == 0 | -0.21123 | 0.01050 | -20.118 | <1e-05 *** |
| plv - aec == 0 | -0.05249 | 0.01050 | -4.999 | <1e-05 *** |
| pli - aec_orth == 0 | 0.26734 | 0.01050 | 25.462 | <1e-05 *** |
| plv - aec_orth == 0 | 0.42608 | 0.01050 | 40.581 | <1e-05 *** |
| plv - pli == 0 | 0.15874 | 0.01050 | 15.119 | <1e-05 *** |

| | Estimate | Std. Error | z-value | Pr(>\|z\|) |
|---|---|---|---|---|
| lcmv - eloreta == 0 | 0.13484 | 0.01050 | 12.842 | < 1e-06 *** |
| wmne - eloreta == 0 | 0.08206 | 0.01050 | 7.815 | < 1e-06 *** |
| wmne - lcmv == 0 | -0.05278 | 0.01050 | -5.027 | 1.51e-06 *** |

| | Estimate | Std. Error | z-value | Pr(>\|z\|) |
|---|---|---|---|---|
| lcmv.aec - eloreta.aec == 0 | 0.134836 | 0.010500 | 12.842 | <0.001 *** |
| wmne.aec - eloreta.aec == 0 | 0.082056 | 0.010500 | 7.815 | <0.001 *** |
| eloreta.aec_orth - eloreta.aec == 0 | -0.478563 | 0.010500 | -45.580 | <0.001 *** |
| lcmv.aec_orth - eloreta.aec == 0 | -0.425745 | 0.010500 | -40.549 | <0.001 *** |
| wmne.aec_orth - eloreta.aec == 0 | -0.529538 | 0.010500 | -50.435 | <0.001 *** |
| eloreta.pli - eloreta.aec == 0 | -0.211226 | 0.010500 | -20.118 | <0.001 *** |
| lcmv.pli - eloreta.aec == 0 | -0.152836 | 0.010500 | -14.556 | <0.001 *** |
| wmne.pli - eloreta.aec == 0 | -0.157184 | 0.010500 | -14.971 | <0.001 *** |
| eloreta.plv - eloreta.aec == 0 | -0.052486 | 0.010500 | -4.999 | <0.001 *** |
| lcmv.plv - eloreta.aec == 0 | 0.182442 | 0.010500 | 17.376 | <0.001 *** |
| wmne.plv - eloreta.aec == 0 | 0.064648 | 0.010500 | 6.157 | <0.001 *** |
| wmne.aec - lcmv.aec == 0 | -0.052780 | 0.010500 | -5.027 | <0.001 *** |
| eloreta.aec_orth - lcmv.aec == 0 | -0.613398 | 0.010500 | -58.422 | <0.001 *** |
| lcmv.aec_orth - lcmv.aec == 0 | -0.560581 | 0.010500 | -53.391 | <0.001 *** |
| wmne.aec_orth - lcmv.aec == 0 | -0.664373 | 0.010500 | -63.277 | <0.001 *** |
| eloreta.pli - lcmv.aec == 0 | -0.346062 | 0.010500 | -32.960 | <0.001 *** |
| lcmv.pli - lcmv.aec == 0 | -0.287672 | 0.010500 | -27.399 | <0.001 *** |
| wmne.pli - lcmv.aec == 0 | -0.292019 | 0.010500 | -27.813 | <0.001 *** |
| eloreta.plv - lcmv.aec == 0 | -0.187322 | 0.010500 | -17.841 | <0.001 *** |
| lcmv.plv - lcmv.aec == 0 | 0.047606 | 0.010500 | 4.534 | <0.001 *** |
| wmne.plv - lcmv.aec == 0 | -0.070188 | 0.010500 | -6.685 | <0.001 *** |
| eloreta.aec_orth - wmne.aec == 0 | -0.560618 | 0.010500 | -53.395 | <0.001 *** |
| lcmv.aec_orth - wmne.aec == 0 | -0.507800 | 0.010500 | -48.364 | <0.001 *** |
| wmne.aec_orth - wmne.aec == 0 | -0.611593 | 0.010500 | -58.250 | <0.001 *** |
| eloreta.pli - wmne.aec == 0 | -0.293281 | 0.010500 | -27.933 | <0.001 *** |
| lcmv.pli - wmne.aec == 0 | -0.234891 | 0.010500 | -22.372 | <0.001 *** |
| wmne.pli - wmne.aec == 0 | -0.239239 | 0.010500 | -22.786 | <0.001 *** |

eloreta.plv - wmne.aec == 0 -0.134542 0.010500 -12.814 <0.001 ***
lcmv.plv - wmne.aec == 0 0.100387 0.010500 9.561 <0.001 ***
wmne.plv - wmne.aec == 0 -0.017408 0.010500 -1.658 0.887
lcmv.aec_orth - eloreta.aec_orth == 0 0.052818 0.010500 5.031 <0.001 ***
wmne.aec_orth - eloreta.aec_orth == 0 -0.050975 0.010500 -4.855 <0.001 ***
eloreta.pli - eloreta.aec_orth == 0 0.267337 0.010500 25.462 <0.001 ***
lcmv.pli - eloreta.aec_orth == 0 0.325727 0.010500 31.023 <0.001 ***
wmne.pli - eloreta.aec_orth == 0 0.321379 0.010500 30.609 <0.001 ***
eloreta.plv - eloreta.aec_orth == 0 0.426076 0.010500 40.581 <0.001 ***
lcmv.plv - eloreta.aec_orth == 0 0.661005 0.010500 62.956 <0.001 ***
wmne.plv - eloreta.aec_orth == 0 0.543211 0.010500 51.737 <0.001 ***
wmne.aec_orth - lcmv.aec_orth == 0 -0.103793 0.010500 -9.886 <0.001 ***
eloreta.pli - lcmv.aec_orth == 0 0.214519 0.010500 20.431 <0.001 ***
lcmv.pli - lcmv.aec_orth == 0 0.272909 0.010500 25.993 <0.001 ***
wmne.pli - lcmv.aec_orth == 0 0.268561 0.010500 25.578 <0.001 ***
eloreta.plv - lcmv.aec_orth == 0 0.373258 0.010500 35.550 <0.001 ***
lcmv.plv - lcmv.aec_orth == 0 0.608187 0.010500 57.925 <0.001 ***
wmne.plv - lcmv.aec_orth == 0 0.490393 0.010500 46.706 <0.001 ***
eloreta.pli - wmne.aec_orth == 0 0.318312 0.010500 30.317 <0.001 ***
lcmv.pli - wmne.aec_orth == 0 0.376702 0.010500 35.878 <0.001 ***
wmne.pli - wmne.aec_orth == 0 0.372354 0.010500 35.464 <0.001 ***
eloreta.plv - wmne.aec_orth == 0 0.477051 0.010500 45.436 <0.001 ***
lcmv.plv - wmne.aec_orth == 0 0.711980 0.010500 67.811 <0.001 ***
wmne.plv - wmne.aec_orth == 0 0.594186 0.010500 56.592 <0.001 ***
lcmv.pli - eloreta.pli == 0 0.058390 0.010500 5.561 <0.001 ***
wmne.pli - eloreta.pli == 0 0.054042 0.010500 5.147 <0.001 ***
eloreta.plv - eloreta.pli == 0 0.158739 0.010500 15.119 <0.001 ***
lcmv.plv - eloreta.pli == 0 0.393668 0.010500 37.494 <0.001 ***
wmne.plv - eloreta.pli == 0 0.275874 0.010500 26.275 <0.001 ***
wmne.pli - lcmv.pli == 0 -0.004348 0.010500 -0.414 1.000
eloreta.plv - lcmv.pli == 0 0.100350 0.010500 9.558 <0.001 ***
lcmv.plv - lcmv.pli == 0 0.335278 0.010500 31.933 <0.001 ***
wmne.plv - lcmv.pli == 0 0.217484 0.010500 20.714 <0.001 ***
eloreta.plv - wmne.pli == 0 0.104697 0.010500 9.972 <0.001 ***
lcmv.plv - wmne.pli == 0 0.339626 0.010500 32.347 <0.001 ***
wmne.plv - wmne.pli == 0 0.221832 0.010500 21.128 <0.001 ***
lcmv.plv - eloreta.plv == 0 0.234929 0.010500 22.375 <0.001 ***
wmne.plv - eloreta.plv == 0 0.117134 0.010500 11.156 <0.001 ***
wmne.plv - lcmv.plv == 0 -0.117794 0.010500 -11.219 <0.001 ***

**DAN - Pearson Correlation**

| Factors | F | Pr(>Chisq) |
|---|---|---|
| Inverse method (2, 4727) | 19649.99 | 2.2e-16 *** |
| Connectivity measure (3, 4727) | 560700.50 | 2.2e-16 *** |
| Number of channels (1, 4727) | 8141.99 | 2.2e-16 *** |
| Inverse method × connectivity measure (6, 4727) | 7864.98 | 2.2e-16 *** |
| Inverse method × channels (2, 4727) | 31.88 | 2.2e-16 *** |
| Connectivity measure × channels (3, 4727) | 536.30 | 2.2e-16 *** |

| | | |
|---|---|---|
| Inverse method × connectivity measure × channels (6, 4727) | 43.48 | 2.2e-16 *** |

R2m = 0.9971
R2c = 0.9973

| | Estimate | Std. Error | z-value | Pr(>\|z\|) |
|---|---|---|---|---|
| aec_orth - aec == 0 | 0.66243 | 0.00141 | 469.8 | <2e-16 *** |
| pli - aec == 0 | 0.59786 | 0.00141 | 424.0 | <2e-16 *** |
| plv - aec == 0 | 0.41127 | 0.00141 | 291.7 | <2e-16 *** |
| pli - aec_orth == 0 | -0.06457 | 0.00141 | -45.8 | <2e-16 *** |
| plv - aec_orth == 0 | -0.25116 | 0.00141 | -178.1 | <2e-16 *** |
| plv - pli == 0 | -0.18659 | 0.00141 | -132.3 | <2e-16 *** |

| | Estimate | Std. Error | z-value | Pr(>\|z\|) |
|---|---|---|---|---|
| lcmv - eloreta == 0 | -0.06275 | 0.00141 | -44.5 | <2e-16 *** |
| wmne - eloreta == 0 | -0.04738 | 0.00141 | -33.6 | <2e-16 *** |
| wmne - lcmv == 0 | 0.01537 | 0.00141 | 10.9 | <2e-16 *** |

| | Estimate | Std. Error | z-value | Pr(>\|z\|) |
|---|---|---|---|---|
| lcmv.aec - eloreta.aec == 0 | -0.035927 | 0.003153 | -11.395 | <0.01 *** |
| wmne.aec - eloreta.aec == 0 | -0.027880 | 0.003153 | -8.842 | <0.01 *** |
| eloreta.aec_orth - eloreta.aec == 0 | 0.694961 | 0.003153 | 220.418 | <0.01 *** |
| lcmv.aec_orth - eloreta.aec == 0 | 0.516339 | 0.003153 | 163.765 | <0.01 *** |
| wmne.aec_orth - eloreta.aec == 0 | 0.792996 | 0.003153 | 251.511 | <0.01 *** |
| eloreta.pli - eloreta.aec == 0 | 0.614961 | 0.003153 | 195.045 | <0.01 *** |
| lcmv.pli - eloreta.aec == 0 | 0.610630 | 0.003153 | 193.671 | <0.01 *** |
| wmne.pli - eloreta.aec == 0 | 0.591876 | 0.003153 | 187.723 | <0.01 *** |
| eloreta.plv - eloreta.aec == 0 | 0.394882 | 0.003153 | 125.243 | <0.01 *** |
| lcmv.plv - eloreta.aec == 0 | 0.273970 | 0.003153 | 86.894 | <0.01 *** |
| wmne.plv - eloreta.aec == 0 | 0.310619 | 0.003153 | 98.518 | <0.01 *** |
| wmne.aec - lcmv.aec == 0 | 0.008047 | 0.003153 | 2.552 | 0.307 |
| eloreta.aec_orth - lcmv.aec == 0 | 0.730888 | 0.003153 | 231.813 | <0.01 *** |
| lcmv.aec_orth - lcmv.aec == 0 | 0.552266 | 0.003153 | 175.160 | <0.01 *** |
| wmne.aec_orth - lcmv.aec == 0 | 0.828923 | 0.003153 | 262.906 | <0.01 *** |
| eloreta.pli - lcmv.aec == 0 | 0.650888 | 0.003153 | 206.440 | <0.01 *** |
| lcmv.pli - lcmv.aec == 0 | 0.646557 | 0.003153 | 205.066 | <0.01 *** |
| wmne.pli - lcmv.aec == 0 | 0.627803 | 0.003153 | 199.118 | <0.01 *** |
| eloreta.plv - lcmv.aec == 0 | 0.430809 | 0.003153 | 136.638 | <0.01 *** |
| lcmv.plv - lcmv.aec == 0 | 0.309897 | 0.003153 | 98.289 | <0.01 *** |
| wmne.plv - lcmv.aec == 0 | 0.346546 | 0.003153 | 109.913 | <0.01 *** |
| eloreta.aec_orth - wmne.aec == 0 | 0.722840 | 0.003153 | 229.260 | <0.01 *** |
| lcmv.aec_orth - wmne.aec == 0 | 0.544218 | 0.003153 | 172.607 | <0.01 *** |
| wmne.aec_orth - wmne.aec == 0 | 0.820876 | 0.003153 | 260.354 | <0.01 *** |
| eloreta.pli - wmne.aec == 0 | 0.642841 | 0.003153 | 203.887 | <0.01 *** |
| lcmv.pli - wmne.aec == 0 | 0.638510 | 0.003153 | 202.513 | <0.01 *** |
| wmne.pli - wmne.aec == 0 | 0.619756 | 0.003153 | 196.565 | <0.01 *** |
| eloreta.plv - wmne.aec == 0 | 0.422762 | 0.003153 | 134.086 | <0.01 *** |
| lcmv.plv - wmne.aec == 0 | 0.301850 | 0.003153 | 95.736 | <0.01 *** |
| wmne.plv - wmne.aec == 0 | 0.338499 | 0.003153 | 107.360 | <0.01 *** |
| lcmv.aec_orth - eloreta.aec_orth == 0 | -0.178622 | 0.003153 | -56.653 | <0.01 *** |
| wmne.aec_orth - eloreta.aec_orth == 0 | 0.098035 | 0.003153 | 31.093 | <0.01 *** |
| eloreta.pli - eloreta.aec_orth == 0 | -0.079999 | 0.003153 | -25.373 | <0.01 *** |
| lcmv.pli - eloreta.aec_orth == 0 | -0.084331 | 0.003153 | -26.747 | <0.01 *** |

| | | | | |
|---|---|---|---|---|
| wmne.pli - eloreta.aec_orth == 0 | -0.103085 | 0.003153 | -32.695 | <0.01 *** |
| eloreta.plv - eloreta.aec_orth == 0 | -0.300079 | 0.003153 | -95.175 | <0.01 *** |
| lcmv.plv - eloreta.aec_orth == 0 | -0.420991 | 0.003153 | -133.524 | <0.01 *** |
| wmne.plv - eloreta.aec_orth == 0 | -0.384342 | 0.003153 | -121.900 | <0.01 *** |
| wmne.aec_orth - lcmv.aec_orth == 0 | 0.276657 | 0.003153 | 87.746 | <0.01 *** |
| eloreta.pli - lcmv.aec_orth == 0 | 0.098622 | 0.003153 | 31.280 | <0.01 *** |
| lcmv.pli - lcmv.aec_orth == 0 | 0.094291 | 0.003153 | 29.906 | <0.01 *** |
| wmne.pli - lcmv.aec_orth == 0 | 0.075537 | 0.003153 | 23.958 | <0.01 *** |
| eloreta.plv - lcmv.aec_orth == 0 | -0.121457 | 0.003153 | -38.522 | <0.01 *** |
| lcmv.plv - lcmv.aec_orth == 0 | -0.242369 | 0.003153 | -76.871 | <0.01 *** |
| wmne.plv - lcmv.aec_orth == 0 | -0.205720 | 0.003153 | -65.247 | <0.01 *** |
| eloreta.pli - wmne.aec_orth == 0 | -0.178035 | 0.003153 | -56.467 | <0.01 *** |
| lcmv.pli - wmne.aec_orth == 0 | -0.182366 | 0.003153 | -57.840 | <0.01 *** |
| wmne.pli - wmne.aec_orth == 0 | -0.201120 | 0.003153 | -63.788 | <0.01 *** |
| eloreta.plv - wmne.aec_orth == 0 | -0.398114 | 0.003153 | -126.268 | <0.01 *** |
| lcmv.plv - wmne.aec_orth == 0 | -0.519026 | 0.003153 | -164.617 | <0.01 *** |
| wmne.plv - wmne.aec_orth == 0 | -0.482377 | 0.003153 | -152.994 | <0.01 *** |
| lcmv.pli - eloreta.pli == 0 | -0.004331 | 0.003153 | -1.374 | 0.969 |
| wmne.pli - eloreta.pli == 0 | -0.023085 | 0.003153 | -7.322 | <0.01 *** |
| eloreta.plv - eloreta.pli == 0 | -0.220079 | 0.003153 | -69.802 | <0.01 *** |
| lcmv.plv - eloreta.pli == 0 | -0.340991 | 0.003153 | -108.151 | <0.01 *** |
| wmne.plv - eloreta.pli == 0 | -0.304342 | 0.003153 | -96.527 | <0.01 *** |
| wmne.pli - lcmv.pli == 0 | -0.018754 | 0.003153 | -5.948 | <0.01 *** |
| eloreta.plv - lcmv.pli == 0 | -0.215748 | 0.003153 | -68.428 | <0.01 *** |
| lcmv.plv - lcmv.pli == 0 | -0.336660 | 0.003153 | -106.777 | <0.01 *** |
| wmne.plv - lcmv.pli == 0 | -0.300011 | 0.003153 | -95.153 | <0.01 *** |
| eloreta.plv - wmne.pli == 0 | -0.196994 | 0.003153 | -62.480 | <0.01 *** |
| lcmv.plv - wmne.pli == 0 | -0.317906 | 0.003153 | -100.829 | <0.01 *** |
| wmne.plv - wmne.pli == 0 | -0.281257 | 0.003153 | -89.205 | <0.01 *** |
| lcmv.plv - eloreta.plv == 0 | -0.120912 | 0.003153 | -38.349 | <0.01 *** |
| wmne.plv - eloreta.plv == 0 | -0.084263 | 0.003153 | -26.725 | <0.01 *** |
| wmne.plv - lcmv.plv == 0 | 0.036649 | 0.003153 | 11.624 | <0.01 *** |

**DAN - Closeness accuracy**

| Factors | F | Pr(>Chisq) |
|---|---|---|
| Inverse method (2, 4727) | 30.04 | 1.083e-13 *** |
| Connectivity measure (3, 4727) | 10254.87 | 2.2e-16 *** |
| Number of channels (1, 4727) | 101.50 | 2.2e-16 *** |
| Inverse method × connectivity measure (6, 4727) | 43.76 | 2.2e-16 *** |
| Inverse method × channels (2, 4727) | 1.05 | 0.3518 |
| Connectivity measure × channels (3, 4727) | 32.53 | 2.2e-16 *** |
| Inverse method × connectivity measure × channels (6, 4727) | 0.96 | 0.4516 |

| | **Estimate** | **Std. Error** | **z-value** | **Pr(>\|z\|)** |
|---|---|---|---|---|
| aec_orth - aec == 0 | -1.02044 | 0.02129 | -47.926 | < 1e-09 *** |
| pli - aec == 0 | -0.88826 | 0.02129 | -41.718 | < 1e-09 *** |
| plv - aec == 0 | 0.15834 | 0.02129 | 7.437 | < 1e-09 *** |
| pli - aec_orth == 0 | 0.13218 | 0.02129 | 6.208 | 1.32e-09 *** |
| plv - aec_orth == 0 | 1.17878 | 0.02129 | 55.362 | < 1e-09 *** |

| | | | | |
|---|---|---|---|---|
| plv - pli == 0 | 1.04660 | 0.02129 | 49.154 | < 1e-09 *** |

| | **Estimate** | **Std. Error** | **z-value** | **Pr(>\|z\|)** |
|---|---|---|---|---|
| lcmv - eloreta == 0 | 0.028906 | 0.021292 | 1.358 | 0.363 |
| wmne - eloreta == 0 | -0.008618 | 0.021292 | -0.405 | 0.914 |
| wmne - lcmv == 0 | -0.037523 | 0.021292 | -1.762 | 0.182 |
